# Synthesis and single-molecule imaging reveal stereospecific enhancement of binding kinetics by the antitumor eEF1A antagonist SR-A3

**DOI:** 10.1101/2020.10.06.325498

**Authors:** Hao-Yuan Wang, Haojun Yang, Mikael Holm, Keely Oltion, Harrison Tom, Amjad Ayad Qatran Al-Khdhairawi, Jean-Frédéric F. Weber, Scott C. Blanchard, Davide Ruggero, Jack Taunton

## Abstract

Ternatin-family cyclic peptides inhibit protein synthesis by targeting the eukaryotic elongation factor-1α (eEF1A). A potentially related cytotoxic natural product (“A3”) was isolated from Aspergillus, but only 4 of its 11 stereocenters could be assigned. Here, we synthesized SR-A3 and SS-A3 – two out of 128 possible A3 epimers – and discovered that synthetic SR-A3 is indistinguishable from naturally derived A3. Relative to SS-A3, SR-A3 exhibits enhanced residence time and rebinding kinetics, as revealed by single-molecule fluorescence imaging of elongation reactions catalyzed by eEF1A in vitro. Increased residence time – stereospecifically conferred by the unique β-hydroxyl in SR-A3 – was also observed in cells. Consistent with its prolonged duration of action, thrice-weekly dosing with SR-A3 led to dramatically increased survival in an aggressive Myc-driven mouse lymphoma model. Our results demonstrate the potential of SR-A3 as a cancer therapeutic and exemplify an evolutionary mechanism for enhancing cyclic peptide binding kinetics via stereospecific side-chain hydroxylation.

## Main

All living organisms rely on protein synthesis mediated by the ribosome and its associated translation factors. Bacterial ribosomes have long been targeted by small-molecule antimicrobials,^1^ while the human ribosome and translation factors have recently emerged as promising drug targets for cancer and viral infections.^2,3^ Eukaryotic elongation factor-1α (eEF1A) is an essential component of the translation machinery.^4^ GTP-bound eEF1A delivers aminoacyl-transfer RNAs (aa-tRNA) to the ribosomal A site during the elongation phase of protein synthesis. Base pairing between the A-site mRNA codon and the aa-tRNA anticodon promotes GTP hydrolysis by eEF1A, releasing the aa-tRNA from eEF1A and allowing its accommodation into the ribosome. The growing protein chain is subsequently transferred from the P-site peptidyl tRNA to the A-site aa-tRNA, extending it by one amino acid through ribosome-catalyzed peptide bond formation.

Tumor cells and viruses highjack the protein synthesis machinery to elicit growth and replication. Specific eEF1A inhibitors – all of which are macrocyclic natural products – have been evaluated as potential anticancer and antiviral drugs.^5^ Didemnin B,^6,7^ cytotrienin A,^8^ nannocystin A,^9^ and cordyheptapeptide A^10^ are examples of structurally diverse macrocycles that bind eEF1A and inhibit translation elongation. Dehydro-didemnin B (plitidepsin) is approved in Australia for the treatment of relapsed/refractory multiple myeloma^11^ and is efficacious in SARS-CoV-2 infection models.^12^

The cyclic heptapeptide A3 was isolated from an Aspergillus strain on the basis of its ability to inhibit cancer cell proliferation at low nanomolar concentrations.^13^ Although the amino acid composition, sequence, and *N*-methylation pattern of A3 were deduced, only 4 out of 11 stereocenters could be assigned (Figure 1). Motivated by its potent antiproliferative activity and unknown mechanism of action, we sought to determine which of the 128 possible stereoisomers (based on 7 unassigned stereocenters) corresponds to A3. Based on our observation that the amino acid sequence and *N*-methylation pattern of A3 and ternatin are similar,^14^ we previously designed and synthesized ternatin-4, which incorporates the dehydromethyl leucine (dhML) and pipecolic acid residues found in A3, yet lacks the β-hydroxy group attached to *N*-Me-Leu (Figure 1). We discovered that ternatin-4 inhibits cancer cell proliferation and SARS-CoV-2 replication by targeting eEF1A.^15,16^ However, the precise step(s) of eEF1A-catalyzed elongation blocked by ternatin-family cyclic peptides – as well as the structure of A3 and the role of its unique β-hydroxy group – all remain unknown.

**Figure 1.**
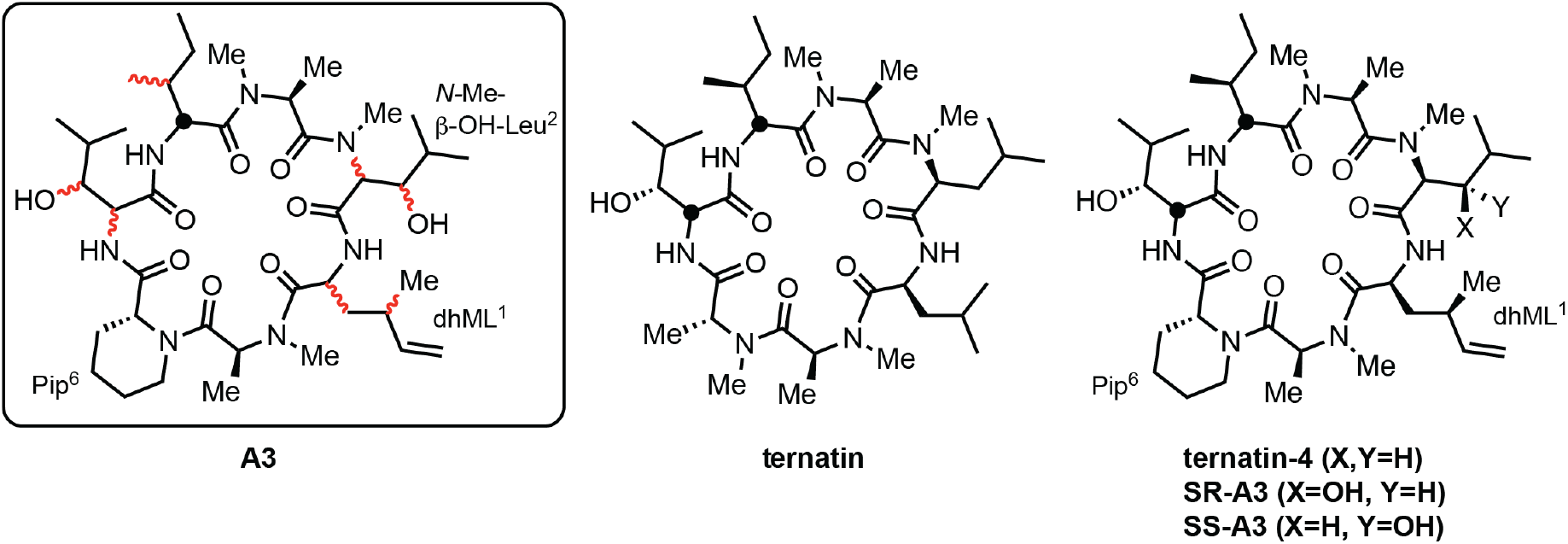
Partially determined structure of the natural product A3. We hypothesized that A3 corresponds to one of two epimers – SR-A3 or SS-A3 – related to ternatin, a natural product, and ternatin-4, an A3-inspired synthetic compound.^14^

Here, we report the first total syntheses of two A3 epimers, SR-A3 and SS-A3 (Figure 1), along with cellular and single-molecule biophysical studies focused on quantifying drug-target residence times. Synthetic SR-A3 potently inhibited cell proliferation and protein synthesis by targeting eEF1A and was spectroscopically and biologically indistinguishable from the natural product A3. Transient exposure of cells to SR-A3 resulted in long-lasting inhibitory effects, whereas similarly prolonged effects were not observed with SS-A3 or ternatin-4. To gain mechanistic insight into these differences, we assessed eEF1A-catalyzed translation via single-molecule fluorescence resonance energy transfer (smFRET) imaging. These experiments directly revealed SR-A3’s comparatively long duration of eEF1A blockade on the ribosome, enabled in part by its enhanced capacity to rebind following dissociation. Preclinical studies in a mouse model of human Burkitt lymphoma revealed that SR-A3 exhibits potent antitumor activity. Our data thus reveal a striking and stereospecific enhancement in eEF1A binding kinetics conferred by a single oxygen atom appended to a cyclic peptide.

## Results and Discussion

### Synthesis of SR-A3 and SS-A3 via an expeditious route to dehydromethyl-Leu

We speculated that dehydromethyl leucine (hereafter “dhML”) in the natural product A3 has the same stereochemistry as in the synthetic compound, ternatin-4 (Figure 1). Because our original 6-step synthesis of dhML methyl ester was low yielding and required a costly chiral auxiliary, we developed a more efficient, second-generation synthesis suitable for preparing gram quantities of Fmoc-dhML.

Copper(I)-promoted S_N_2’ reaction of a serine-derived organozinc reagent with allylic electrophiles has been previously used to synthesize amino acids that contain a γ-stereogenic center.^17,18^ This method was appealing because it would provide dhML (as the Boc methyl ester) in only two steps from the inexpensive chiral building block, Boc-(*S*)-serine-OMe.^17^ After extensive optimization aimed at improving S_N_2’ vs. S_N_2 selectivity and conversion (Supplementary Figure 1), we obtained Boc-dhML-OMe **3** in 43% isolated yield (1.6 g) through the use of 50 mol% CuBrDMS and 2 equivalents of crotyl chloride (Figure 2a). Boc to Fmoc exchange, followed by ester hydrolysis, provided Fmoc-dhML **5**, which was incorporated into the linear heptapeptide as described below.

**Figure 2.**
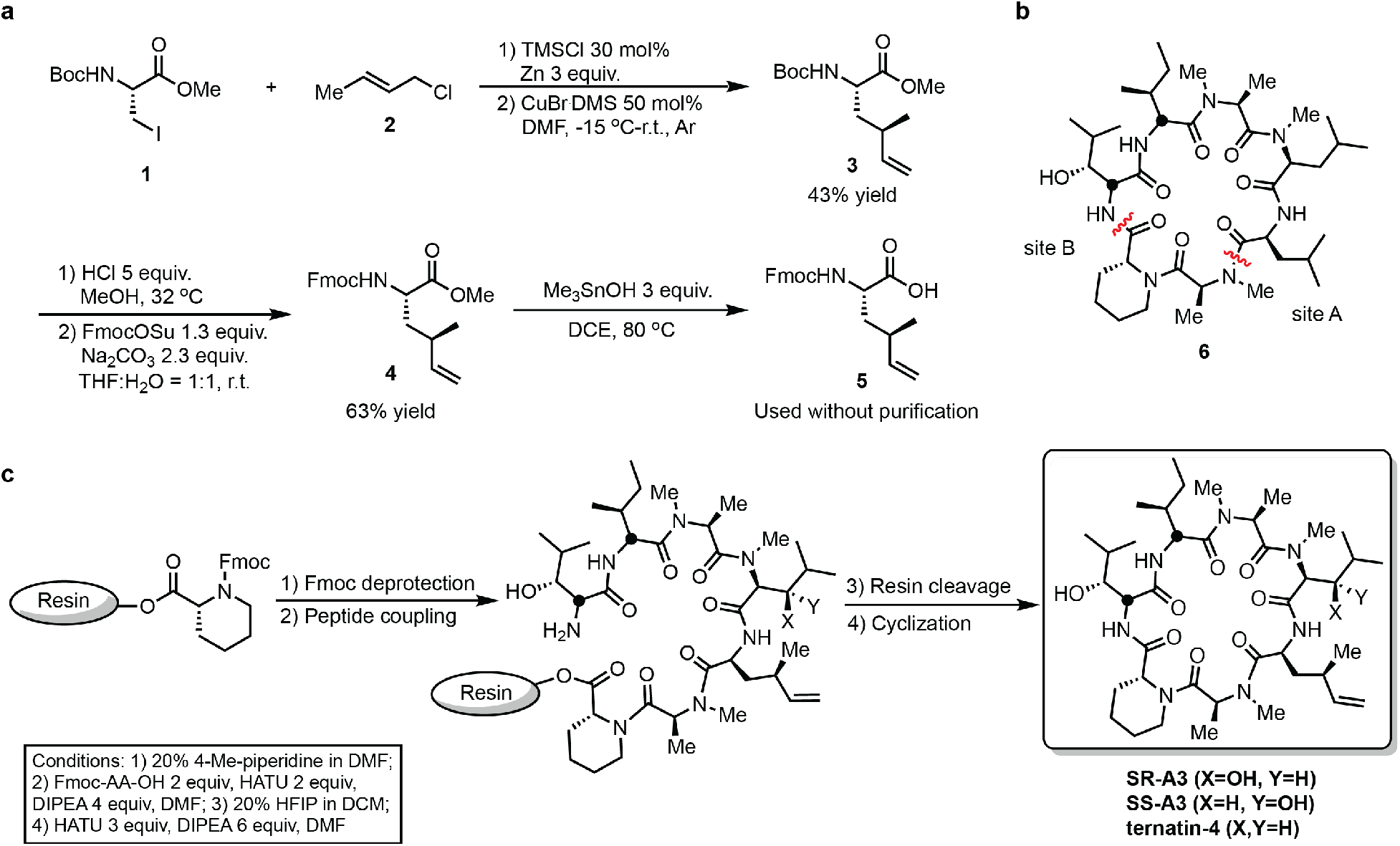
Expeditious synthesis of dhML, ternatin-4, and A3 epimers. (**a**) Scheme for synthesis of Fmoc-dhML **5**. (**b**) Identification of alternative macrocyclization site B. (**c**) Scheme for solid-phase synthesis of linear heptapeptide precursors, followed by solution-phase cyclization to provide ternatin-4, SR-A3, and SS-A3 (see Supporting Information for details).

A solid-phase route was previously employed to synthesize a linear heptapeptide precursor of ternatin, followed by solution-phase cyclization.^14^ However, this strategy involved macrocyclization between the secondary amine of *N*-Me-Ala7 and the carboxylic acid of Leu1 (Figure 2b, site A), which we found to be low yielding in the context of peptides containing dhML.^15^ Thus, we sought to identify an alternative cyclization site using the ternatin-related cyclic peptide **6** as a model system (Figure 2c). Linear heptapeptide precursors were synthesized on the solid phase, deprotected and cleaved from the resin, and cyclized in solution (see Supporting Information for details). Gratifyingly, cyclization at site B provided **6** in 63% overall yield (including the solid-phase linear heptapeptide synthesis), whereas cyclization at site A was less efficient (46% overall yield). By synthesizing the linear heptapeptide precursor on the solid phase and cyclizing in solution at site B, we were able to prepare ternatin-4 in 3 days and 70% overall yield (27 mg), a significant improvement over our previous route (Figure 2c). Most importantly, by incorporating Fmoc-protected (*S,R*)- and (*S,S*)-*N*-Me-β-OH-Leu, we completed the first total syntheses of SR-A3 (21 mg, 35% overall yield) and SS-A3 (5 mg, 21% overall yield).

### Synthetic SR-A3 is indistinguishable from naturally derived A3

With synthetic SR-A3 and SS-A3 in hand (Figure 3a), we first compared their HPLC elution profiles with an authentic sample of the Aspergillus-derived natural product A3. SR-A3 and naturally derived A3 had identical retention times, whereas SS-A3 eluted later in the gradient (Figure 3b). Furthermore, the ^1^H and ^13^C NMR spectra of SR-A3 were identical to the corresponding spectra of natural A3 (Figure 3c). Finally, SR-A3 and naturally derived A3 blocked proliferation of HCT116 cancer cells with superimposable dose-response curves (Figure 3d, IC_50_ ~0.9 nM), whereas SS-A3 was ~3-fold less potent (IC_50_ ~2.7 nM). Similar trends were observed in four additional cancer cell lines (Supplementary Figure 2). Together, these data are consistent with our stereochemical hypothesis and strongly suggest that synthetic SR-A3 and natural A3 have the same structure.

**Figure 3.**
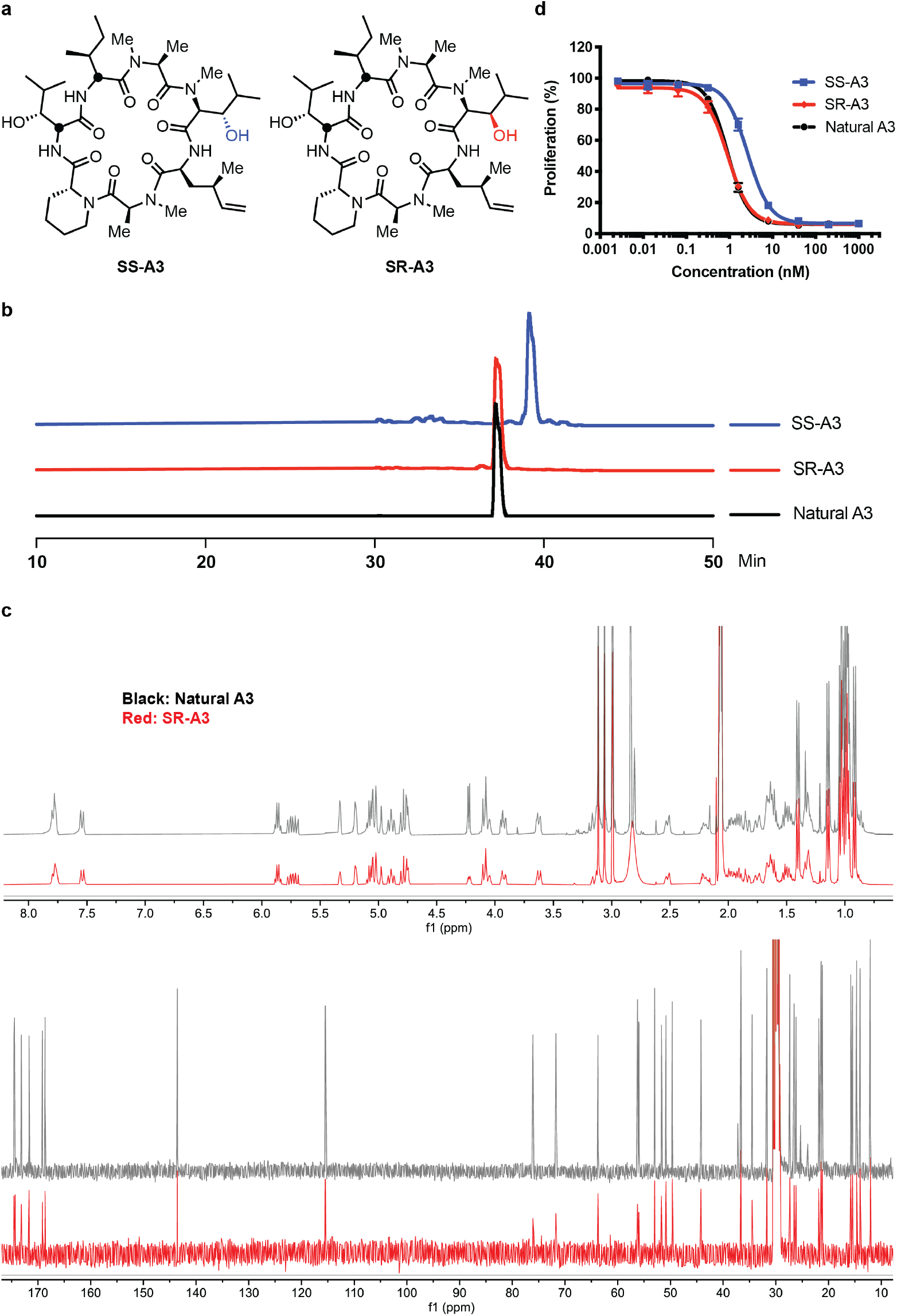
SR-A3 is indistinguishable from naturally derived A3. (**a**) Chemical structures of SS-A3 and SR-A3. (**b**) HPLC elution profiles. (**c**) Overlay of ^1^H and ^13^C NMR spectra in acetone-d_6_. (**d**) Concentration-dependent antiproliferative effects in HCT116 cells after 72 h. Cell proliferation (% DMSO control) was quantified using alamarBlue. Data points (% DMSO control) are mean values ± SD (n = 3).

### *N*-Me-β-OH-Leu stereospecifically confers increased cellular residence time

We previously demonstrated that ternatin-4 is inactive toward a human cancer cell line that is homozygous for an Ala399Val mutation in eEF1A (encoded by the *EEF1A1* gene).^15^ These cells were also resistant to SR-A3 (IC_50_ >> 1 μM), providing strong genetic evidence that eEF1A is the relevant target (Figure 4a). Consistent with this interpretation, treatment of cells with SR-A3 for 24 h inhibited global protein synthesis with an IC_50_ of ~20 nM (Figure 4b), as measured by a clickable puromycin (*O*-propargyl puromycin, OPP) incorporation assay (Supplementary Figure 3).^19^ Under these conditions – 24 h of continuous treatment prior to a 1-h pulse with OPP — SR-A3 behaved identically to ternatin-4, whereas SS-A3 was slightly less potent. Based on these cellular data, it remained unclear as to whether *N*-Me-β-OH-Leu in SR-A3 confers any advantage over the biosynthetically less ornate *N*-Me-Leu found in ternatin and ternatin-4.

**Figure 4.**
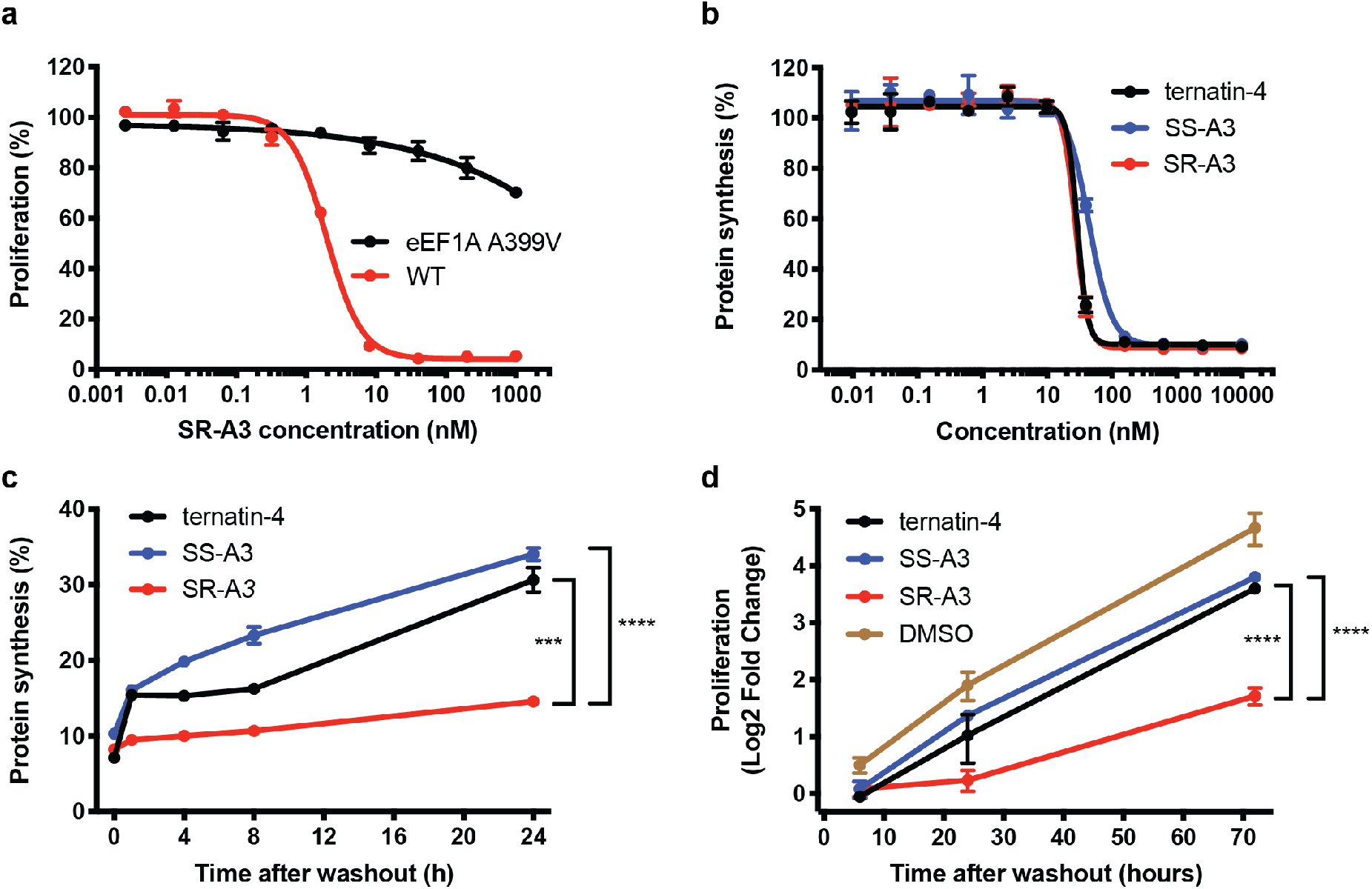
*N*-Me-β-OH-Leu stereospecifically endows SR-A3 with increased cellular residence time. (**a**) Wild-type and eEF1A-mutant (A399V) HCT116 cells were treated with SR-A3 for 72 h. Cell proliferation (% DMSO control) was quantified using alamarBlue. (**b**) HCT116 cells were treated with the indicated compounds for 24 h and protein synthesis was quantified after pulse labeling with *O*-propargyl puromycin for 1 h (see Supporting Information). Data points (% DMSO control) are mean values ± SD (n = 3). (**c**) HCT116 cells were treated with the indicated compounds (100 nM) or DMSO for 4 h, followed by washout into compound-free media. At the indicated time points post-washout, cells were pulse-labeled with OPP (1 h), and OPP incorporation was quantified. Normalized data (% DMSO control) are mean values ±SD (n = 3). (**d**) HCT116 cells were treated with the indicated compounds (100 nM) or DMSO for 4 h, followed by washout into compound-free media. At the indicated time points post-washout, cell proliferation was quantified using the CellTiter-Glo assay. Normalized data (log2 fold change vs. DMSO control at t = 0 h post-washout) are mean values ±SD (n = 3). ***, P < 0.001; ****, P < 0.0001.

Drug-target residence time, which reflects not only the biochemical off-rate, but also the rebinding rate and local target density in vivo, has emerged as a critical kinetic parameter in drug discovery.^20,21^ To test for potential differences in cellular residence time, we treated HCT116 cells with 100 nM SR-A3, SS-A3, or ternatin-4 for 4 h, followed by washout into drug-free media. At various times post-washout, cells were pulse-labeled with OPP for 1 h. Whereas protein synthesis rates partially recovered in cells treated with ternatin-4 or SS-A3 (~30% of DMSO control levels, 24 h post-washout), transient exposure of cells to SR-A3 resulted in more prolonged inhibition (Figure 4c). To confirm the extended duration of action observed with SR-A3, we assessed cell proliferation during a 72-h washout period. Strikingly, cell proliferation over 72 h was sharply reduced after 4-h treatment with 100 nM SR-A3, followed by rigorous washout. By contrast, cell proliferation rates recovered nearly to DMSO control levels after transient exposure to 100 nM ternatin-4 or SS-A3 (Figure 4d). These results demonstrate that the (*R*)-β-hydroxy group attached to *N*-Me-Leu endows SR-A3 with a substantial kinetic advantage over SS-A3 and ternatin-4, as reflected by washout resistance and increased cellular residence time associated with inhibition of protein synthesis and cell proliferation.

### smFRET imaging reveals enhanced residence time and rebinding kinetics of SR-A3

To gain mechanistic insight into SR-A3’s kinetic advantage in cells relative to ternatin-4 and SS-A3, we employed smFRET imaging of biochemically reconstituted, eEF1A-catalyzed translation reactions.^22^ In this assay, tRNA in the P site of surface-immobilized human ribosomes is labeled with a donor fluorophore (Cy3) (Figure 5a). Aminoacyl-tRNA (aa-tRNA) labeled with an acceptor fluorophore (LD655) is then stopped-flow delivered as a ternary complex (TC) with eEF1A and GTP. The resulting reaction is monitored in real time using a home-built total internal reflection microscope.^23^ The FRET efficiency between the two fluorophores – inversely related to the distance between the two tRNAs – increases through a series of well-characterized states (**S1**-**S4**, Figure 5a).^22,24^

**Figure 5.**
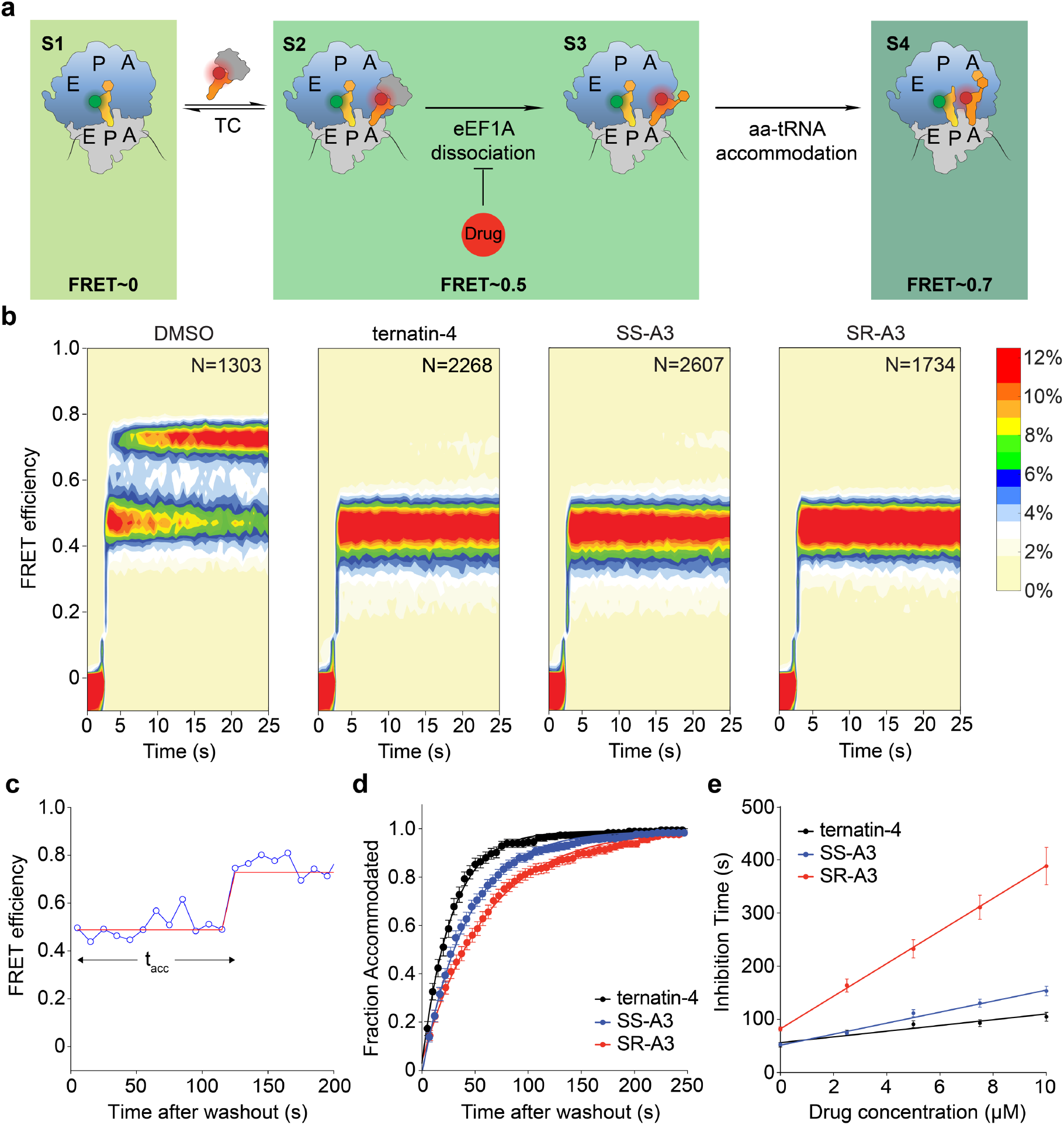
Single-molecule FRET imaging reveals increased residence time and rebinding kinetics of SR-A3. (**a**) Schematic showing predominant long-lived reaction species (**S1**-**S4**) detected by the experimental setup. (**b**) Population FRET histograms of reactions initiated from **S1** by delivering the ternary complex (TC) and either DMSO or the indicated drug (10 μM) at the start of data acquisition. N = number of observed molecules. (**c**) Representative smFRET trace from a chase experiment initiated from drug-stalled elongation complexes (**S2**) by pre-incubation with SR-A3, followed by washout into drug-free buffer at the start of data acquisition. Blue circles represent the measured FRET efficiency for each movie frame; the solid red line represents a Hidden-Markov model idealization of the data. (**d**) Cumulative drug dissociation time distributions constructed from FRET traces exemplified by (**c**) after washout into drug-free buffer. Data points represent bootstrap means (± SEM) from all FRET traces acquired in a single movie. The solid lines represent fits of single-exponential functions to the data. (**e**) Plots of inhibition time (prior to accommodation) as a function of drug concentration in the washout buffer were used to determine drug residence times and rebinding constants. Inhibition times were derived from curves exemplified in (**d**), after washout into buffer containing the indicated drug concentrations. Each data point represents the mean of two experimental replicates (± SEM). Solid lines are linear fits to the data, adjusted R^2^ = 0.90 (ternatin-4), 0.98 (SS-A3), and 0.99 (SR-A3).

Before TC addition, no FRET is observed (Figure 5a, **S1**; Figure 5b). When TC is introduced to the microscope flow cell, it binds rapidly to the ribosomal A site where codon-anticodon recognition occurs (Figure 5a, **S2**), resulting in GTP hydrolysis and subsequent dissociation of eEF1A from aa-tRNA and the ribosome (Figure 5a, **S3**). Within both of these states (Figure 5a, **S2** and **S3**), the ribosome complex exhibits similar time-averaged FRET efficiencies of ~0.5. Upon eEF1A dissociation, aa-tRNA accommodates into the ribosomal peptidyl transferase center (Figure 5a, **S4**), as revealed by the marked increase in FRET efficiency to ~0.7 in DMSO control reactions (Figure 5b, left). By contrast, when TC was delivered in the presence of 10 μM ternatin-4, SS-A3, or SR-A3, aa-tRNA accommodation was strongly inhibited, whereas formation of the initial TC/ribosome intermediate was unaffected (Figure 5b). These results suggest that all three compounds similarly stall elongating ribosomes bound to TC (Figure 5a, **S2**), presumably by preventing conformational changes in eEF1A (either before or after GTP hydrolysis), which are required for its dissociation from aa-tRNA and the ribosome.

Since SR-A3 exhibited greater cellular residence time than ternatin-4 or SS-A3 (Figure 4e), we designed a series of in vitro chase experiments to quantify differences in dissociation rates and rebinding constants between these closely related analogs. We first generated a population of drug-stalled elongation complexes by incubating immobilized ribosomes with TC and 10 μM of each drug for 30 seconds. Excess TC and drugs were then washed out of the microscope flow cell with buffer containing 0, 2.5, 5, 7.5, or 10 μM drug at the start of data acquisition. Under these washout conditions, drug-stalled elongation complexes can gradually transition into the fully accommodated high-FRET state (Figure 5c, example trace showing washout into drug-free buffer). We estimated the average time required for aa-tRNA accommodation to occur in each washout condition (t_acc_) from cumulative dwell-time distributions by calculating the fraction of ribosomes that reached the high-FRET state at each time point and fitting exponential functions to the resulting data (Figure 5d and Supplementary Figure 4a, see Supporting Information for details).

Given the assumption that drug dissociation must occur prior to aa-tRNA accommodation, the drug residence times and rebinding constants can then be determined from the chase series (Figure 5e, see Supporting Information for details).^25,26^ Based on the measured inhibition time after washout into drug-free buffer, the residence time of SR-A3 (82 s) is about 60% longer than ternatin-4 (56 s) or SS-A3 (51 s). In addition, the rebinding constant, which corresponds to the drug concentration in the washout buffer required to double the inhibition time (relative to drug-free buffer) through drug rebinding events, strongly favors SR-A3 (Figure 5e). This implies that SR-A3 rebinds to the stalled eEF1A-ribosome complex (**S2** or related states, Figure 5a) twice as fast as SS-A3 and four times as fast as ternatin-4 (Figure 5e and Supplementary Figure 4b). Taken together, and consistent with our cell-based findings, our smFRET data show that (*S,R*)-*N*-Me-β-OH-Leu stereospecifically endows SR-A3 with a longer residence time and faster target rebinding kinetics upon dissociation.

### SR-A3 treatment significantly extends survival of Eμ-Myc tumor-bearing mice

The oncogenic transcription factor Myc is dysregulated in >50% of human cancers.^27^ Structural alterations of the *MYC* gene cause B-cell lymphoma in humans and mice,^28–31^ and a Myc-dependent increase in protein synthesis is a key oncogenic determinant.^32^ The Eμ-Myc transgenic mouse,^28^ in which Myc is specifically overexpressed in B lymphocytes, has been employed as a preclinical model of human Burkitt lymphoma and other Myc-driven B-cell malignancies. Hence, the Eμ-Myc model is a paradigm for testing whether inhibition of protein synthesis downstream of Myc oncogenic activity confers therapeutic benefit. Given its enhanced residence time and rebinding kinetics, as well as improved metabolic stability (Supplementary Figure 5), we selected SR-A3 for a preclinical trial using the Eμ-Myc lymphoma allograft model. After intravenous injection of wild-type immunocompetent mice with mouse Eμ–Myc/+ lymphoma cells and waiting until tumors were palpable (~2 weeks after injection), we began treatment with either vehicle or SR-A3 (dosed 3 times per week, 1.5 or 2.0 mg/kg, by intraperitoneal injection). Treatment of tumor-bearing mice with single-agent SR-A3 dramatically prolonged survival in a dose-dependent manner (Figure 6). Moreover, SR-A3 was well tolerated in both dose groups, and no significant body weight loss was observed (Supplementary Figure 6). These results suggest that SR-A3, a next-generation eEF1A antagonist, is a viable preclinical candidate for the treatment of Myc-driven B lymphoid tumors.

**Figure 6.**
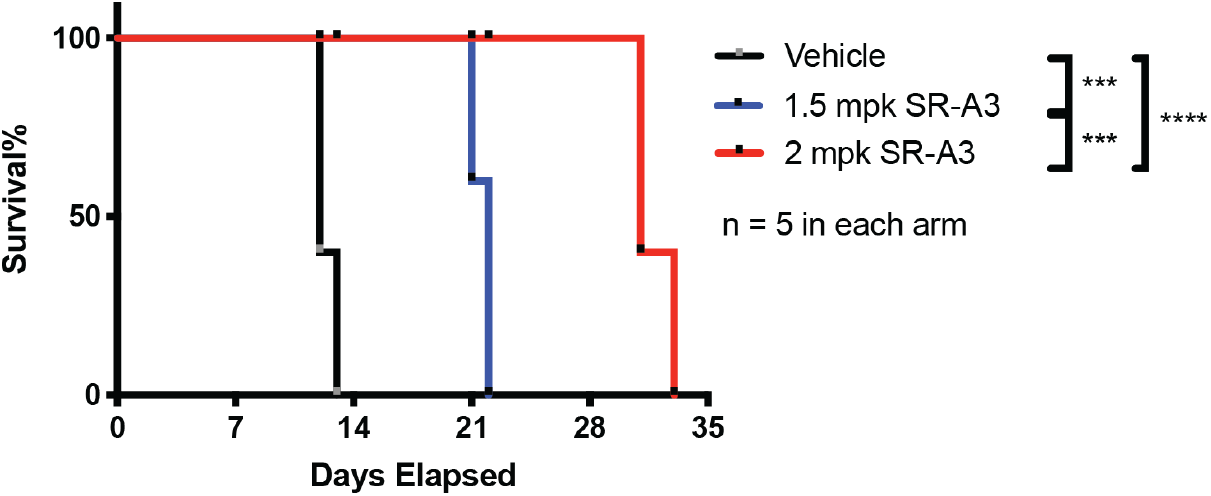
SR-A3 extends overall survival of Eμ-Myc tumor-bearing mice. Kaplan-Meier survival curves for mice dosed with vehicle or the indicated doses of SR-A3 (intraperitoneal injection, 3 times per week). Day 0 indicates the beginning of treatment. ****, P < 0.0001, ***, P < 0.001.

## Conclusions

In this study, we first developed an improved synthetic route to dhML-containing ternatin variants, culminating in the first total syntheses of SR-A3 and SS-A3 (Figure 2). Our work provides spectroscopic, chromatographic, and pharmacological evidence that synthetic SR-A3 (and not SS-A3) is identical to the fungal natural product “A3” (Figure 3), confirming the previous partial structure elucidation and providing the first complete stereochemical assignment of this potent eEF1A antagonist.

SR-A3 differs structurally from the previously reported eEF1A inhibitor, ternatin-4, by the addition of a single oxygen atom into the side chain of *N*-Me-Leu (Figure 1). We speculate that A3 is evolutionarily related to ternatin via acquisition of biosynthetic modules for (*R*)-pipecolic acid and (*S*,*R*)-dhML, as well as stereospecific hydroxylation of *N*-Me-Leu by the A3-producing fungus. Although amino acid β-hydroxylation is a common biosynthetic modification in cyclic peptide natural products, its stereospecific functions are mostly unknown. An unexpected finding from our work is that the *N*-Me-Leu β-hydroxyl in SR-A3 has little effect on cellular potency under continuous treatment conditions, as compared with ternatin-4. Rather, the β-hydroxyl in SR-A3, but not SS-A3, confers a dramatic increase in drug-target residence time and rebinding kinetics, as revealed by washout experiments in cells and reconstituted eEF1A-catalyzed elongation reactions monitored by smFRET (Figure 4d,e and Figure 5). We note that “drug-target residence time” measured in cells and tissues is determined by both drug dissociation and rebinding kinetics. Moreover, the intracellular concentration (or local density) of the target can play a major role in drug rebinding kinetics.^20^ This can potentially explain the much longer residence time in cells, where [eEF1A] is ~35 μM,^33^ as compared to the smFRET flow cell, where [eEF1A] is <<10 nM post-washout. The structural basis of SR-A3’s enhanced binding kinetics will likely require cryo-electron microscopy analysis of stalled SR-A3/eEF1A/ribosome complexes at atomic resolution. Although the precise molecular mechanism by which the β-hydroxyl confers this kinetic advantage awaits further investigation, our study nevertheless reveals the power of smFRET imaging to illuminate differences in drug dissociation and rebinding kinetics, both of which can contribute to drug-target residence time and therapeutic efficacy.^20^

Dysregulation of the transcription factor Myc underlies multiple human cancers.^27^ Nevertheless, Myc is still considered “undruggable” and no direct Myc inhibitors have advanced into clinical trials. Because the oncogenic activity of Myc relies on its ability to promote ribosome biogenesis and protein synthesis, an alternative approach is to attenuate protein synthesis rates. The Eμ-Myc lymphoma model has been employed to test various protein synthesis inhibition strategies, often in combination with other cytotoxic drugs. Targeting the protein synthesis machinery in Eμ-Myc mice either genetically^32^ or pharmacologically^34–36^ has been shown to suppress tumor growth and prolong overall survival. However, to the best of our knowledge, no translation elongation inhibitors have shown single-agent efficacy in this aggressive B-cell lymphoma model.

Our preclinical trial revealed that intermittent, low doses of SR-A3 (1.5-2.0 mg/kg, three times per week) profoundly extended the survival of Eμ-Myc mice without obvious toxicity (Figure 6). SR-A3 thus shows therapeutic potential for the treatment of Myc-driven B-cell lymphoma. Our study of SR-A3 chemistry and biology provides a compelling illustration of how a “ligand efficient” side-chain modification (one oxygen atom) can be exploited to alter the pharmacological properties and residence time of a cyclic peptide natural product.

## Data availability

Data supporting the findings of this study are available within the article and the Supplementary Information. Source data are provided with this paper.

## Acknowledgments

Funding for this study was provided by the UCSF Program for Breakthrough Biomedical Research (J.T. and D.R.), the UCSF Invent Fund (J.T. and D.R.), the National Institutes of Health (5R01GM079238 to S.C.B. and R35CA242986 to D.R.), the American Cancer Society (American Cancer Society Research Professor Award to D.R.), and the Tobacco-Related Disease Research Program Postdoctoral Fellowship Awards (28FT-0014 to H.Y.W.). Part of this work was supported by Taylor’s University PhD Scholarship program (A.A.Q.A.K.), as well as a research grant from the Ministry of Education of Malaysia FRGS [600-IRMI/FRGS 5/3 (011/2017) to J.F.W.].

