## Supplementary Information for "Synthesis and single-molecule imaging reveal stereospecific enhancement of binding kinetics by the antitumor eEF1A antagonist SR-A3"

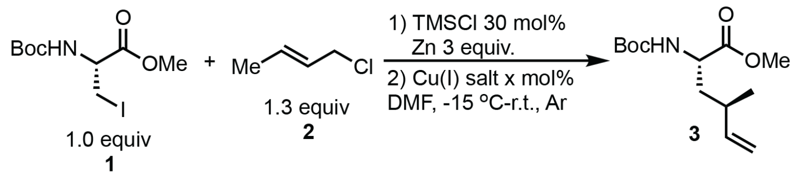


| Entry | Cu(I) Salt | x | $S_N2'$:$S_N2$ [a] | d.r. of $S_N2'$ [a] | Yield [b] |
|---|---|---|---|---|---|
| 1 | CuBr·DMS | 10 | 1.2:1 | 1:1 | 12% |
| 2 | CuBr·DMS | 50 | 1.9:1 | 1.4:1 | 26% |
| 3 | CuBr·DMS | 100 | 2.1:1 | 1.4:1 | - |
| 4 | CuBr | 100 | 1:3.5 | - | - |
| 5 | CuCl | 100 | 1:2.5 | - | - |
| 6 | CuTC | 100 | - | - | N.R. |
| 7[c] | CuBr·DMS | 50 | 1:2.6 | - | - |
| **8[d]** | **CuBr·DMS** | **50** | **1.8:1** | **1.4:1** | **43%** |

[a] Ratio was based on the crude NMR of the reaction; [b] Isolated yield of **3**; [c] Crotyl bromide was used instead of **2**; [d] 2 equiv of crotyl chloride were used. DMS: dimethyl sulfide; TC: thiophene -2-carboxylate; N.R.: no reaction.

**Supplementary Figure 1 (related to Figure 2).** Screening conditions to synthesize Boc-dhML-OMe **3** via Cu(I)-promoted $S_N2'$ reaction.

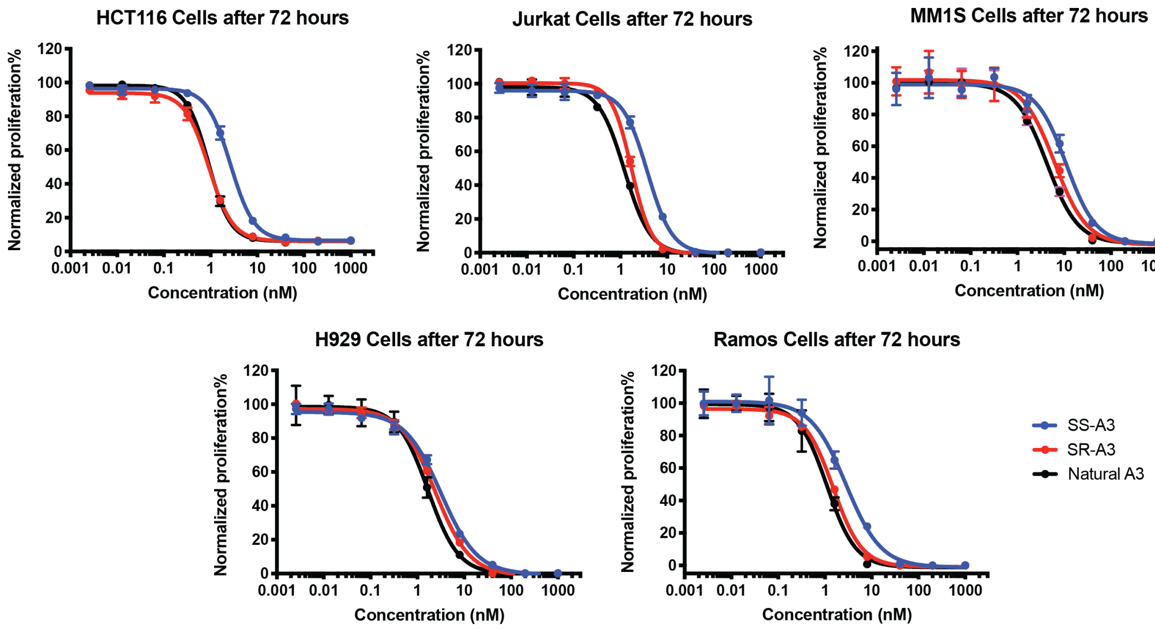


**Supplementary Figure 2 (related to Figure 3).** Antiproliferative effects of SS-A3, SR-A3, and A3 (five-fold dilutions) in cancer cell lines after 72 hours treatment. Cell proliferation (% DMSO control) was quantified using alamarBlue. Data points (% DMSO control) are mean values ± SD (n = 3).

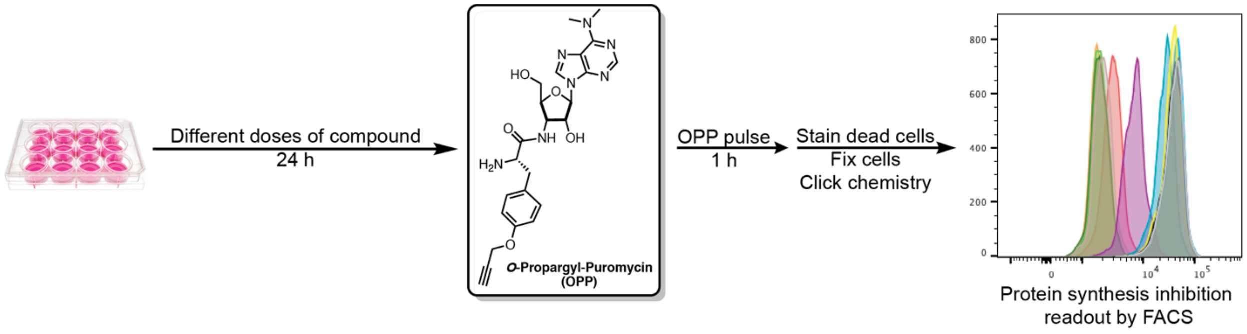


**Supplementary Figure 3 (related to Figure 4).** General workflow for measuring protein synthesis rates in cells using *O*-Propargyl-Puromycin (OPP).

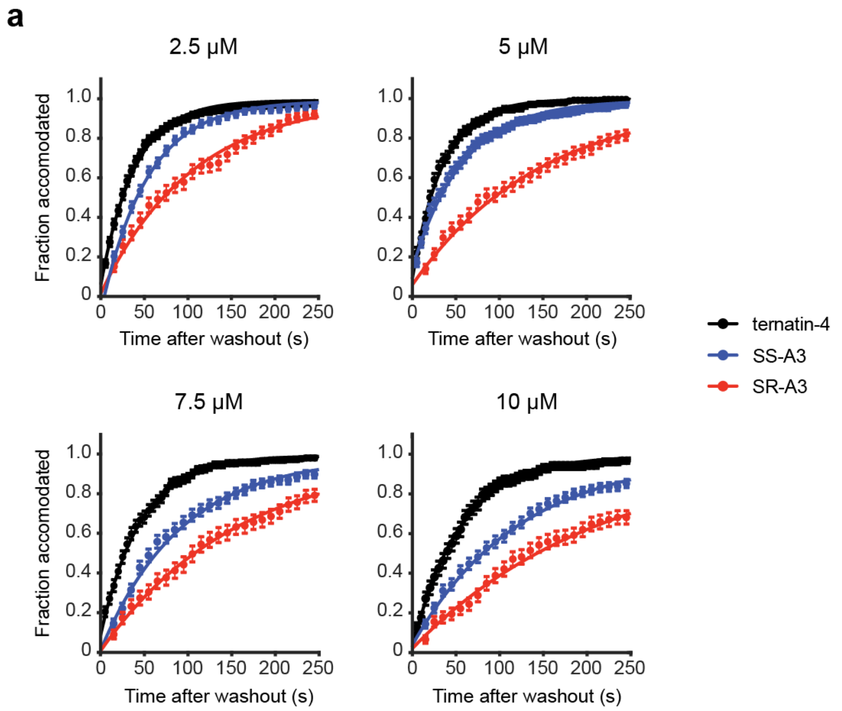


**b**

| | Residence Time (s) | Rebinding Constant (µM) |
|---|---|---|
| ternatin-4 | 56 ± 4 | 10.4 ± 1.9 |
| SS-A3 | 51 ± 2.5 | 5 ± 0.4 |
| SR-A3 | 82 ± 1 | 2.67 ± 0.05 |

**Supplementary Figure 4 (related to Figure 5).** (**a**) Cumulative dissociation time distributions for ternatin-4, SS-A3, and SR-A3 at the indicated concentrations. Distributions are constructed as described in the main text and the methods. Error bars represent SEM derived from 1000 bootstrap replicates. (**b**) Tabulated kinetic parameters based on data in Figure 5 and Supplementary Figure 4a (see smFRET analysis methods below).

| | Human Liver Microsomes (remaining%) | Mouse Liver Microsomes (remaining%) |
|---|---|---|
| ternatin-4 | 1.6% | 1.2% |
| SSA3 | 6.7% | 7.4% |
| SRA3 | 39.9% | 31.8% |

**Supplementary Figure 5 (related to Figure 6).** Human and mouse liver microsome stability results. Percent remaining of each analog was quantified by LC/MS after incubating at 1 μM in the presence of human or mouse liver microsomes (with NADPH) for 30 min at 37°C. This study was performed by the contract research organization, Bioduro-Sundia (San Diego, CA).

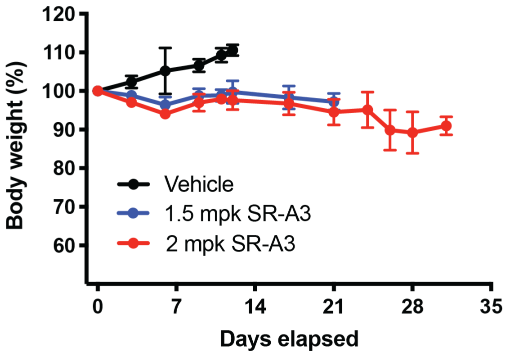


**Supplementary Figure 6.** Average body weight of each group (± SD, relative to day 0) during the SR-A3 efficacy study in Eμ-Myc mice. Day 0 indicates the beginning of treatment.

### Cell culture

HCT116 cells (ATCC, Manassas, VA) were maintained in McCoy's 5A media (Gibco, Grand Island, NY) supplemented with 10% fetal bovine serum (Axenia Biologix, Dixon, CA), 100 units/mL penicillin, and 100 µg/mL streptomycin (Gibco). H929 cells (ATCC) were maintained in advanced RPMI 1640 media (Gibco) supplemented with 6% fetal bovine serum, 2 mM glutamine, 100 units/mL penicillin, and 100 mg/mL streptomycin. MM1S, Jurkat, and Ramos cells (ATCC) were maintained in RPMI 1640 media (Gibco) supplemented with 10% fetal bovine serum, 100 units/mL penicillin, and 100 mg/mL streptomycin. All cells were cultured at 37°C in a 5% $CO_2$ atmosphere.

### Proliferation assay

Adherent cells were briefly trypsinized and repeatedly pipetted to produce a homogeneous cell suspension. 2,500 cells were seeded in 100 µL complete growth media per well in 96-well clear-bottom plates. Suspension cells were repeatedly pipetted to produce a homogenous cell suspension. 10,000 cells were seeded in 100 µL complete growth media per well in 96-well clear-bottom plates. After allowing cells to grow/adhere overnight, cells were treated with 25 µL/well 5x drug stocks (0.1% DMSO final) and incubated for 72 hours. AlamarBlue (Life Technologies, Grand Island, NY) was used to assess cell viability per the manufacturer's instructions. Briefly, 12.5 µL alamarBlue reagent was added to each well, and plates were incubated at 37°C. Fluorescence intensity was measured every 30 min to determine the linear range for each assay (Ex 545 nm, Em 590 nm, SPARK, Tecan Austria GmbH, Austria). Proliferation curves were generated by first normalizing fluorescence intensity in each well to the DMSO-treated plate average. Normalized fluorescence intensity was plotted in GraphPad Prism (GraphPad, La Jolla, CA), and $IC_{50}$ values were calculated from nonlinear regression curves. The reported $IC_{50}$ values represent the average of at least three independent determinations (± SD).

### Washout-proliferation assay

Adherent cells were briefly trypsinized and repeatedly pipetted to produce a homogenous cell suspension. 2,500 cells were seeded in 100 µL complete growth media per well in 96-well clear-bottom plates. After allowing cells to grow/adhere overnight, cells were treated with ternatin analogs (100 nM, 0.1% DMSO final) and incubated for the indicated times. Growth media was carefully removed, cells were washed with warm PBS twice (2x short wash), followed by 5 min incubation in warm media at 37°C (long wash). This "short-long" washing cycle was repeated 3 times. After indicated times post-washout, CellTiter Glo (Promega, Madison, WI) was used to assess cell viability per the manufacturer's instructions. Briefly, after adding 100 µL CellTiter Glo reagent to each well, the plate was rocked at room temperature for 5-10 min, and the luminescence intensity was measured. Proliferation curves were generated by first normalizing luminescence intensity in each well to the average values from the t = 0 time point. Normalized luminescence intensity was plotted in GraphPad Prism (GraphPad, La Jolla, CA). The reported values represent the average of at least three independent determinations (± SD).

### OPP incorporation assay

HCT116 cells at 60% confluency in 12-well plates were incubated with the indicated concentrations of ternatin analogs for 10 min or 24 h at 37°C. After the indicated times, *O*-propargyl-puromycin (30 µM final concentration) was added, and the cells were incubated for 1 hour at 37°C. Subsequently, media was removed, and the cells were trypsinized, collected, and washed twice with ice-cold PBS before transferring to a 96-well V bottom plate. 100 µL Zombie Red (BioLegend, San Diego, CA) solution was added to each well and incubated for 30 min at

RT in the dark. Cells were then washed with 2% FBS in PBS before fixation with 200 µL of 4% PFA in PBS for 15 min on ice in the dark. After washing the cells with 2% FBS in PBS, 200 µL permeabilization buffer (3% FBS, 0.1% saponin in PBS) was added to each well, and the cells were incubated for 5 min at RT in the dark. Cells were then washed and resuspended in 25 µL permeabilization buffer. 100 µL click chemistry mix (50 mM HEPES pH 7.5, 150 mM NaCl, 400 µM TCEP, 250 µM TBTA, 5 µM CF405M-Azide (Biotium, Fremont, CA), 200 µM $CuSO_4$) was added to each well, and cells were incubated at RT in the dark. After overnight incubation, the cells were washed with permeabilization buffer followed by FACS buffer (2% FBS, 1% P/S, 2 mM EDTA, in PBS w/o Ca/Mg). Cells were then resuspended in 200 µL FACS buffer and filtered before FACS analysis (CytoFLEX, Beckman-Coulter, Brea, CA). Protein synthesis inhibition curves were generated by gating for single live cells and plotting mean fluorescence intensity (MFI) relative to the DMSO control values using GraphPad Prism (GraphPad, La Jolla, CA). $IC_{50}$ values were calculated from nonlinear regression curves. The reported values represent the average of at least three independent determinations (± SD).

#### Washout-OPP assay

HCT116 cells at 60% confluency in 12-well plates were incubated with compounds at 100 nM for 4 h at 37°C. Media was carefully removed, and cells were washed with warm PBS twice (2x short wash), followed by 5 min incubation in warm media at 37°C (long wash). After repeating the "short-long" washout cycle 3 times, cells were then resuspended in warm media and incubated at 37°C. After the indicated times post-washout, *O*-propargyl-puromycin (30 µM final concentration) was added, and the cells were incubated for an additional 1 h at 37°C. The media was removed, and the cells were trypsinized, collected, and washed twice with ice-cold PBS before transferring to a 96-well V bottom plate. 100 µL Zombie Red (BioLegend, San Diego, CA) solution was added to each well and incubated for 30 min at RT in the dark. Cells were then washed with 2% FBS in PBS before fixation with 200 µL of 4% PFA in PBS for 15 min on ice in the dark. After washing the cells with 2% FBS in PBS, 200 µL permeabilization buffer (3% FBS, 0.1% saponin in PBS) was added to each well, and cells were incubated for 5 min at RT in the dark. Cells were then washed and resuspended in 25 µL permeabilization buffer. 100 µL click chemistry mix (50 mM HEPES pH 7.5, 150 mM NaCl, 400 µM TCEP, 250 µM TBTA, 5 µM CF405M-Azide (Biotium, Fremont, CA), 200 µM $CuSO_4$) was added to each well, and cells were incubated at RT in the dark. After the overnight incubation, cells were washed with permeabilization buffer followed by FACS buffer (2% FBS, 1% P/S, 2 mM EDTA, in PBS w/o Ca/Mg). Cells were then resuspended in 200 µL FACS buffer and filtered before FACS analysis as described above.

#### smFRET data collection

Ribosomes from HEK293T cells, elongation factor eEF1A, and fluorescence-labeled tRNAs were prepared using the protocol described previously.[1] All smFRET experiments were carried out at 25 °C in human polymix buffer (20 mM HEPES pH 7.5, 5 mM $MgCl_2$, 140 mM KCl, 10 mM $NH_4Cl$, 2 mM spermidine, 5 mM putrescine and 1.5 mM 2-mercaptoethanol) containing 500 µM cycloheximide and a mixture of triplet-state quenchers (1 mM trolox, 1 mM 4-nitrobenzyl alcohol (NBA), 1 mM cyclooctatetraene (COT)) and an enzymatic oxygen scavenging system (2 µM 3,4-dihydroxybenzoic acid (PCA), 0.02 Units/ml protocatechuate *3,4*-dioxygenase (PCD)). The time-evolution of the FRET signal was recorded using a home-built total internal reflection-based fluorescence microscope at ~0.1 kW/$cm^2$ laser (532 nm) illumination. Movies were recorded either in time-lapse mode with one 500 ms frame acquired every 10 seconds or continuously at a time resolution of 500 ms. Donor and acceptor fluorescence intensities were extracted from the recorded movies and FRET efficiency traces were calculated using the SPARTAN software package.[2] FRET traces were selected for further analysis according to the following criteria: a single catastrophic photobleaching event, at least 8:1 signal/background-noise ratio and 6:1 signal/signal-noise ratio, less than four donor-fluorophore blinking events, and a correlation coefficient between donor and acceptor < 0.5. The resulting smFRET traces were analyzed using Hidden Markov model idealization methods as implemented in the SPARTAN software package.[2]

#### Ternary complex real-time delivery experiments

80S initiation complexes containing Met-tRNA$^{fMet}$-Cy3 and displaying the codon UUC in the A site were immobilized on passivated quartz slides as described previously.[1,3] This was followed by delivery of 10 nM eEF1A ternary complex containing Phe-tRNA$^{Phe}$-LD655 together with 1 mM GTP and either DMSO or 10 µM drug at the start of data acquisition.

#### Drug chase experiments

Stalled pre-accommodation complexes were formed by immobilization of 80S initiation complexes containing Met-tRNA$^{fMet}$-Cy3 and displaying the codon UUC in the A site to passivated quartz slides as described previously[1,3] followed by delivery of 10 nM eEF1A ternary complex containing Phe-tRNA$^{Phe}$-LD655 together with 1 mM GTP and 10 µM drug. After 30 s of incubation, the chase was started by delivery of buffer containing either 0, 2.5, 5, 7.5, or 10 µM drug concurrent with the start of data acquisition.

#### smFRET data kinetic analysis

To construct cumulative distribution plots suitable for estimation of kinetic parameters, FRET traces were first idealized using Hidden Markov model analysis to a model with three FRET states (FRET efficiencies: -0.0043 ± 0.06, 0.4485 ± 0.06, and 0.7070 ± 0.06) and then the cumulative sum of molecules that had arrived at the highest (0.7070) FRET state in each movie frame was calculated. To estimate reaction mean times and their associated uncertainties, 1000 bootstrap samples were generated from each experimental replicate and mean times were estimated by fitting of single-exponential functions to these data (Equation 1), taking into account unobserved events due to photobleaching of the fluorophores or dissociation of the intact ternary complex from the ribosomal A site by multiplying the estimated mean time with the inverse of the fraction of traces were accommodation was observed at the end of the process.[4]

$$f_{acc}(t) = \left(1 - e^{\frac{-1}{\frac{1}{f_{acc}(\infty)}\tau_{acc}}t}\right) \qquad \text{Equation 1}$$

Mean rates and standard errors were calculated as the weighted averages of two to three experimental replicates for each drug concentration. For estimation of drug residence times and rebinding constants these estimated mean times were plotted against drug concentration and Equation 2 was fitted to the data.[4,5]

$$\tau_I([Drug]) = \tau_0 \left(1 + \frac{[Drug]}{K_I}\right) \qquad \text{Equation 2}$$

In Equation 2, $\tau_I$ is the observed inhibition mean time as a function of drug concentration. $\tau_0$ is the drug residence time each time it binds, and $K_I$ is the rebinding constant corresponding to the drug concentration required to double the inhibition time through drug rebinding events, it can be interpreted as the ratio between the drug association rate constant and the rate of the process that renders the ribosome immune to drug inhibition, in this case conformational changes in eEF1A.

**Animal Experiments**

All animal experiments were approved by The University of California San Francisco Institutional Animal Care and Use Committee (UCSF-IACUC). The *Eμ-Myc/+* transgenic mice were purchased from the Jackson Laboratory (stock no. 002728). Primers used for genotyping are: 5'-CCG AGG TGA GTG TGA GAG G-3'; 5'-AAA CAG TAA TAG CGC AGC A-3'.
For *Eμ-Myc* clonal B-cell line collection, *Eμ-Myc/+* mice harboring lymphoma were euthanized according to IACUC guidelines. Lymph nodes were collected immediately on ice, minced, and passed through a 40-μm cell strainer in cold PBS with 2% FBS. Cells were centrifuged at 300 *g* for 5 min. Cells were then resuspended in cold erythrocyte lysis solution ACK (Thermo Fisher A1049201) for 1 min. Isolated lymphoma cells were centrifuged at 300 *g* for 5 min and washed in PBS before freezing in cell cryopreservation medium and storing in liquid nitrogen. For the lymphoma preclinical trial, *Eμ-Myc/+* lymphoma cells were thawed and washed once in PBS. One million cells were injected intravenously into eight-week-old male C57BL/6 mice. Mice were monitored for lymphoma development by palpation every other day. Once the lymphoma tumors became palpable (~2 weeks after tumor cell injection), mice (5 per group) were dosed by intraperitoneal injection with either SR-A3 or vehicle (10% EtOH/Kolliphor EL in water) three times per week (every other day) until the end point.

The natural product A3 was purified as described previously.[6]

**Chemical synthesis (general)**

All reactions in non-aqueous media were conducted under a positive pressure of dry argon in glassware that had been dried in oven prior to use, unless noted otherwise. Anhydrous solutions of reaction mixtures were transferred via an oven-dried syringe or cannula. All solvents were dried prior to use unless noted otherwise. Thin layer chromatography was performed using precoated silica gel plates (EMD Chemical Inc. 60, F254). Flash column chromatography was performed on CombiFlash Rf 200i system (Teledyne Isco, Lincoln, NE). $^1$H and $^{13}$C nuclear magnetic resonance spectra (NMR) were obtained on a Varian (Palo Alto, CA) Inova 400 MHz spectrometer recorded in ppm (δ) downfield of TMS (δ = 0) in $CDCl_3$ unless noted otherwise. Signal splitting patterns were described as singlet (s), doublet (d), triplet (t), quartet (q), quintet (quint), or multiplet (m), with coupling constants (*J*) in hertz. High resolution mass spectra (HRMS) were performed on Waters Xevo G2-XS QToF LC-MS system, eluting with a water/MeCN (+0.1% formic acid) gradient at 0.6 mL/min.

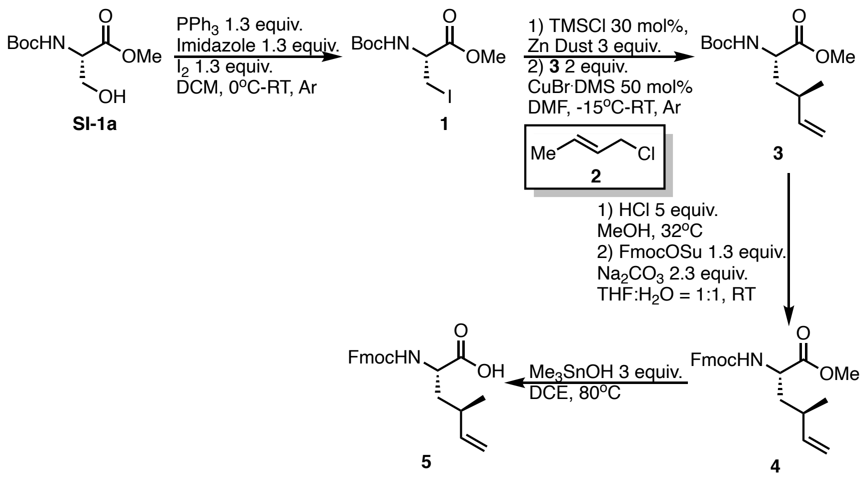

**Synthesis of 5:**

To an oven-dried flask was added $PPh_3$ (59.3 mmol, 15.6 g), imidazole (59.3 mmol, 4.1 g), and anhydrous DCM (250 mL) under Ar. After cooling the mixture to 0 °C, $I_2$ (59.3 mmol, 15.1 g) was added in three portions and the reaction was warmed up to RT and stirred for 10 min. The reaction was then re-cooled to 0 °C, Boc-Ser-OMe **SI-1a** (45.6 mmol, 10 g) was added. After stirring at 0 °C for 1 h, the reaction was allowed to warm up to RT. After completion (~2-3 h), the reaction was filtered through a pad of Celite and concentrated *in vacuo*. To the above filtrate, $Et_2O$ was added to crush out triphenyl phosphite, and the resulting slurry was filtered through a pad of Celite and concentrated *in vacuo*. CombiFlash chromatography (0-10% EA in Hexane) was used to afford **1** as the product (yellow solid, 12.1 g, 81% yield). Spectral data of **1** are in accordance with the literature.[7]

To an oven-dried flask was added Zn dust (44.4 mmol, 2.9 g), anhydrous DMF (20 mL), and TMSCl (4.44 mmol, 0.56 mL) at RT under Ar. After stirring for 30 min, the slurry was cooled to 0 °C and **1** (14.8 mmol, 4.87 g) was added. The mixture was allowed to warm up to RT and stirred for 1 h to form the corresponding Zn reagent. To a separated oven-dried flask was added CuBr·DMS (7.4 mmol, 1.52 g), **2** (29.6 mmol, 2.9 mL), and anhydrous DMF (20 mL) at RT under Ar. After the mixture was cooled to -15 °C, the Zn reagent was added. The reaction was then warmed up to RT and stirred overnight. After completion, the reaction was quenched with $NH_4Cl$ (aq), extracted by EA, dried over $Na_2SO_4$, and concentrated *in vacuo*. CombiFlash chromatography (0-5% EA in Hexane) was used to afford **3** as the product (colorless liquid, 1.64 g, 43% yield). Spectral data of **3** were in accordance with our previous report.[8]

To an oven-dried flask was added **3** (8 mmol, 2.07 g) and 2 M HCl in MeOH (40 mmol, 20 mL). The reaction was stirred at 32 °C for 2 h. After completion, the mixture was concentrated *in vacuo* to afford a crude amine mixture, which was used in the next step without further purification. To an oven-dried flask was added the crude amine mixture, Fmoc-OSu (10.4 mmol, 3.5 g), $Na_2CO_3$ (18.4 mmol, 1.97 g), THF (20 mL), and $H_2O$ (20 mL). The reaction was stirred overnight at RT. After completion, the reaction was quenched with $NH_4Cl$ (aq), extracted by EA, dried over $Na_2SO_4$, and concentrated *in vacuo*. CombiFlash chromatography (0-10% EA in Hexane) was used to afford **4** as the product (colorless liquid, 1.92 g, 63% yield over 2 steps). HRMS (ESI) for $C_{23}H_{25}NNaO_4$ $[M+Na]^+$, 402.1676 (Calc.), found 402.1716.

To an oven-dried flask was added **4** (0.2 mmol, 76 mg), $Me_3SnOH$ (0.6 mmol, 108.5 mg), and DCE (2 mL). The reaction was stirred at 80 °C for 4 h. After completion, the reaction was diluted with EA, washed with 1 M HCl and brine, dried over $Na_2SO_4$, and concentrated *in vacuo*. The crude mixture of **5** was used in the solid phase synthesis without further purification.

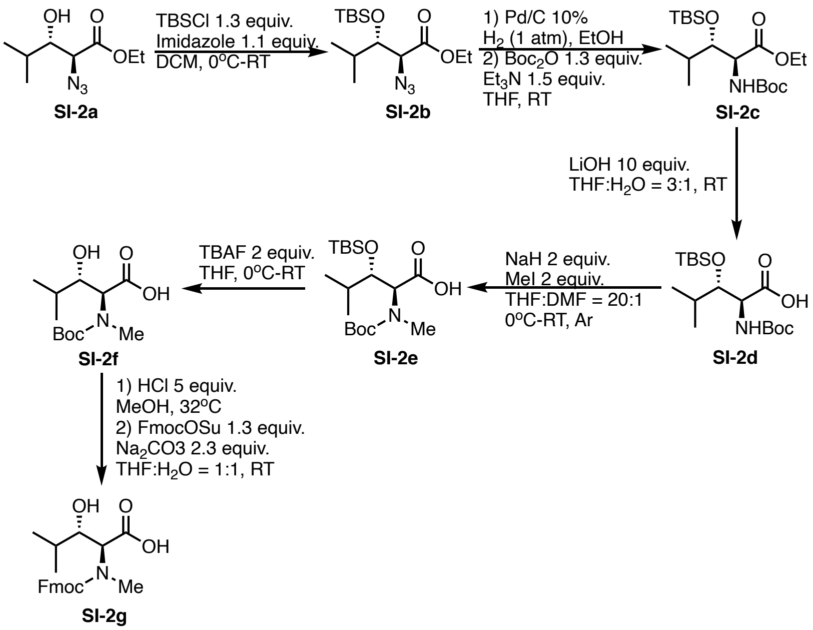

**Synthesis of SI-2g:**

Compound **SI-2a** was synthesized according to the previously reported protocol.[9]

To an oven-dried flask was added **SI-2a** (10 mmol, 2 g), imidazole (13 mmol, 885 mg), and anhydrous DCM (50 mL). After cooling to 0 °C, TBSCl (13 mmol, 1.96 g) was added and the reaction was warmed up to RT and stirred overnight. After completion, the reaction was quenched with $NH_4Cl$ (aq), extracted by DCM, dried over $Na_2SO_4$, and concentrated *in vacuo*. The crude mixture was used in the next step without further purification. To an oven-dried flask was added **SI-2b** crude mixture, Pd/C (10%, 320 mg), and EtOH (50 mL) under $H_2$. The reaction was stirred overnight at RT. After completion, the reaction was filtered through a pad of Celite and concentrated *in vacuo* to afford a crude mixture. To an oven-dried flask was added the above crude mixture, $Boc_2O$ (13 mmol, 3 mL), $Et_3N$ (15 mmol, 2.1 mL) and THF (50 mL). The reaction stirred overnight at RT. After completion, the reaction was quenched with $NH_4Cl$ (aq), extracted by DCM, dried over $Na_2SO_4$, and concentrated *in vacuo*. CombiFlash chromatography (0-10% EA in Hexane) was used to afford **SI-2c** as the product (yellow liquid, 3.2 g, 83% yield after 3 steps). HRMS (ESI) for $C_{19}H_{40}NO_5Si$ $[M+H]^+$, 390.2670 (Calc.), found 390.2664.

To an oven-dried flask was added **SI-2c** (5 mmol, 1.95 g), LiOH (50 mmol, 1.2 g), THF (30 mL), and $H_2O$ (10 mL). The reaction was stirred at RT overnight. After completion, the reaction was quenched with 1 M HCl until pH = 4 and concentrated *in vacuo* to afford a crude mixture, which was used in the next step without further purification. To an oven-dried flask was added **SI-2d** crude mixture, anhydrous THF (40 mL), and anhydrous DMF (2 mL). After the reaction was cooled to 0 °C, NaH (60%, 10 mmol, 400 mg) was added and the reaction was stirred for 30 min at 0 °C. MeI (10 mmol, 0.62 mL) was then added at 0 °C and the reaction was allowed to warm up to RT and stirred overnight. After completion, the reaction was quenched with $NH_4Cl$ (aq), extracted by EA, dried over $Na_2SO_4$, and concentrated *in vacuo*. CombiFlash chromatography (0-10% EA in Hexane) was used to afford **SI-2e** as the product (yellow liquid, 1.24 g, 62% yield after 2 steps). HRMS (ESI) for $C_{18}H_{36}NO_5Si$ $[M-H]^-$, 374.2368 (Calc.), found 374.2409.

To an oven-dried flask was added **SI-2e** (3 mmol, 1.21 g) and THF (30 mL). After the reaction was cooled to 0 °C, TBAF in THF solution (1M, 6 mmol, 6 mL) was added and the reaction was warmed to RT and stirred overnight. After completion, the reaction was quenched with $NH_4Cl$ (aq), extracted by EA, dried over $Na_2SO_4$, and concentrated *in vacuo*. CombiFlash chromatography (0-50% EA with 1% AcOH in Hexane) was used to afford **SI-2f** as the product (yellow liquid, 510 mg, 65% yield). HRMS (ESI) for $C_{12}H_{22}NO_5$ $[M-H]^-$, 260.1498 (Calc.), found 260.1504.

To an oven-dried flask was added **SI-2f** (1 mmol, 261 mg) and 2 M HCl in MeOH (5 mmol, 2.5 mL). The reaction was stirred at 32 °C for 2 h. After completion, the mixture was concentrated *in vacuo* to afford a crude mixture, which was used in the next step without further purification. To an oven-dried flask was added the crude mixture, Fmoc-OSu (1.3 mmol, 439 mg), $Na_2CO_3$ (2.3 mmol, 244 mg), THF (5 mL), and $H_2O$ (5 mL). The reaction was stirred at RT overnight. After completion, the reaction was quenched with $NH_4Cl$ (aq), extracted by EA, dried over $Na_2SO_4$, and concentrated *in vacuo*. CombiFlash chromatography (0-10% MeOH in DCM) was used to afford **SI-2g** as the product (white solid, 238 mg, 62% yield over 2 steps). HRMS (ESI) for $C_{22}H_{24}NO_5$ $[M-H]^-$, 382.1660 (Calc.), found 382.1658.

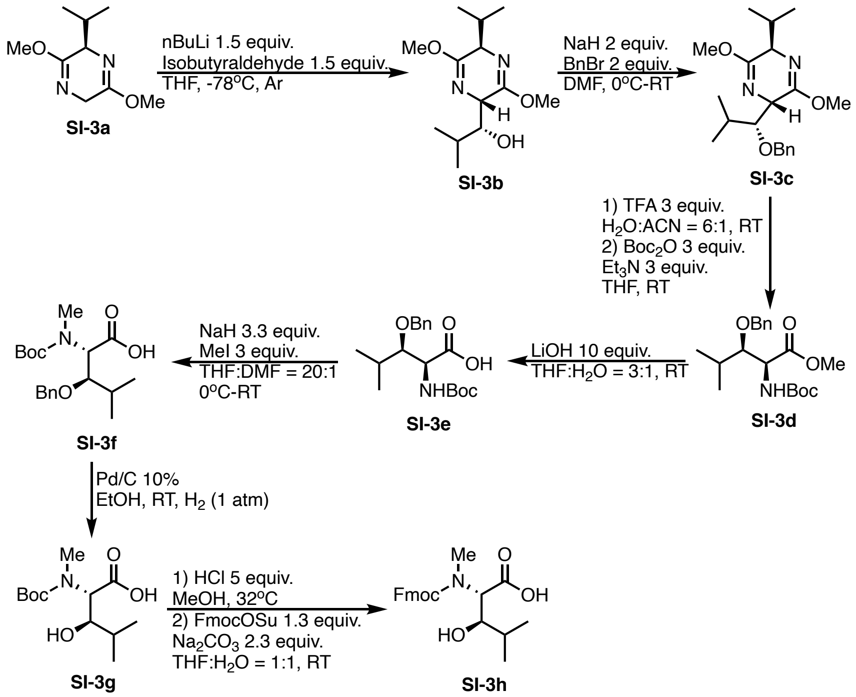


### Synthesis of SI-3h:

To an oven-dried flask was added **SI-3a** (10 mmol, 1.9 g) and anhydrous THF (50 mL) under Ar. After cooling to -78 °C, nBuLi (2.5 M in Hexane, 15 mmol, 6 mL) was added. After stirring at -78 °C for 30 min, isobutyraldehyde (15 mmol, 1.4 mL) was added and the reaction was allowed to warm up to RT and stirred overnight. After completion, the reaction was quenched with $NH_4Cl$ (aq), extracted by DCM, dried over $Na_2SO_4$, and concentrated *in vacuo*. CombiFlash chromatography (0-20% EA in Hexane) was used to afford **SI-3b** as the product (yellow liquid, 1.57 g, 61% yield). Spectral data of **SI-3b** are in accordance with the literature.[10]

To an oven-dried flask was added **SI-3b** (6 mmol, 1.52 g) and anhydrous DMF (50 mL). After cooling to 0 °C, NaH (60%, 12 mmol, 480 mg) was added and the reaction was stirred at 0 °C for 30 min. BnBr (12 mmol, 1.42 mL) was then added and the reaction was allowed to warm up to RT and stirred overnight. After completion, the reaction was quenched with $NH_4Cl$ (aq), extracted by EA, dried over $Na_2SO_4$, and concentrated *in vacuo*. CombiFlash chromatography

(0-10% EA in Hexane) was used to afford **SI-3c** as the product (yellow liquid, 1.98 g, 95% yield). HRMS (ESI) for $C_{20}H_{31}N_2O_3$ $[M+H]^+$, 347.2329 (Calc.), found 347.2341.

To an oven-dried flask was added **SI-3c** (6 mmol, 1.98 g), MeCN (8 mL), and water (48 mL). TFA (18 mmol, 1.4 mL) was added dropwise and the reaction was stirred overnight. After completion, the reaction was concentrated *in vacuo*. The crude mixture was used in the next step without further purification. To an oven-dried flask was added above crude mixture, $Boc_2O$ (18 mmol, 4.2 mL), $Et_3N$ (18 mmol, 2.5 mL), and THF (50 mL). The reaction was stirred overnight at RT. After completion, the reaction was quenched with $NH_4Cl$ (aq), extracted by EA, dried over $Na_2SO_4$, and concentrated *in vacuo*. CombiFlash chromatography (0-10% EA in Hexane) was used to afford **SI-3d** as the product (yellow liquid, 1.33 g, 63% yield after 2 steps). HRMS (ESI) for $C_{19}H_{29}NNaO_5$ $[M+Na]^+$, 374.1938 (Calc.), found 374.1935.

To an oven-dried flask was added **SI-3d** (4 mmol, 1.41 g), LiOH (40 mmol, 1.68 g), THF (30 mL) and water (10 mL). The reaction was stirred at RT for 5 h. After completion, the reaction was concentrated *in vacuo*. The crude mixture was used in the next step without further purification. To an oven-dried flask was added above **SI-3e** crude mixture, anhydrous THF (40 mL) and anhydrous DMF (2 mL). After cooling to 0 °C, NaH (60%, 13.2 mmol, 528 mg) was added. After stirring at 0 °C for 30 min, MeI (12 mmol, 0.75 mL) was added. The reaction was then allowed to warm up to RT and stirred overnight. After completion, the reaction was quenched with $NH_4Cl$ (aq), extracted by EA, dried over $Na_2SO_4$, and concentrated *in vacuo*. CombiFlash chromatography (0-30% EA (with 1% AcOH) in Hexane) was used to afford **SI-3f** as the product (yellow liquid, 745 mg, 53% yield). HRMS (ESI) for $C_{19}H_{28}NO_5$ $[M-H]^-$, 350.1973 (Calc.), found 350.2015.

To an oven-dried flask was added **SI-3f** (2 mmol, 703 mg), Pd/C (10 wt%, ~100 mg), and EtOH (20 mL). The reaction was vigorously stirred at RT under $H_2$ overnight. After completion, the reaction was filtered through a pad of Celite and concentrated *in vacuo*. CombiFlash chromatography (0-50% EA (with 1% AcOH) in Hexane) was used to afford **SI-3g** as the product (yellow liquid, 345 mg, 66% yield). HRMS (ESI) for $C_{12}H_{22}NO_5$ $[M-H]^-$, 260.1498 (Calc.), found 260.1537.

To an oven-dried flask was added **SI-3g** (1 mmol, 261 mg) and 2 M HCl in MeOH (5 mmol, 2.5 mL). The reaction was stirred at 32 °C for 2 h. After completion, the mixture was concentrated *in vacuo*. The crude mixture was used in the next step without further purification. To an oven-dried flask was added the crude mixture, Fmoc-OSu (1.3 mmol, 439 mg), $Na_2CO_3$ (2.3 mmol, 244 mg), THF (5 mL), and $H_2O$ (5 mL) at RT. The reaction was stirred overnight and monitored by TLC. After completion, the reaction was quenched with $NH_4Cl$ (aq), extracted by EA, dried over $Na_2SO_4$, and concentrated *in vacuo*. CombiFlash chromatography (0-10% MeOH in DCM) was used to afford **SI-3h** as the product (white solid, 333 mg, 87% yield over 2 steps). HRMS (ESI) for $C_{22}H_{24}NO_5$ $[M-H]^-$, 382.1660 (Calc.), found 382.1658.

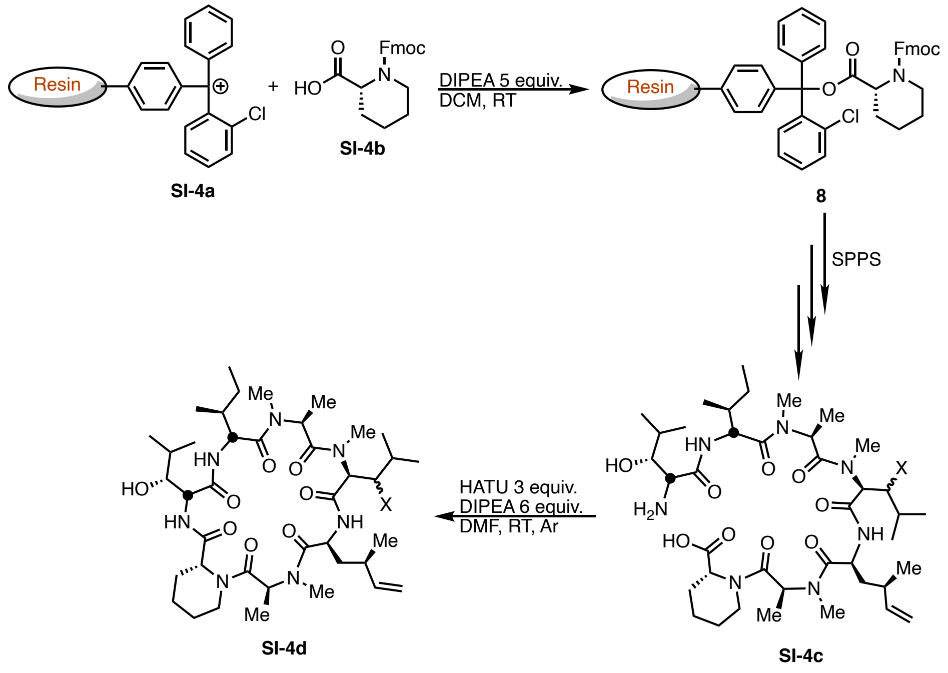

### General procedure for cyclic peptide synthesis:

To a 50 mL syringe with a valve tip was added resin **SI-4a** (Loading = 1.6 mmol/g, 1 mmol, 625 mg) and DCM (30 mL). The syringe was rocked for 1 hour at RT. After 1 h, DCM was filtered out through the valve, and the swelled resin was washed with DCM (10 mL) for three times. **SI-4b** (2 mmol, 708.2 mg), DCM (25 mL), and DIPEA (5 mmol) were then added. After rocking the syringe overnight at RT, liquid was filtered out through the valve, and the resin was washed with DCM (10 mL) for three times. After washing, a mixture of DCM:MeOH:DIPEA = 17:2:1 solution (20 mL) was added into the syringe and agitated for another 1 h. After repeating the previous agitation step again, the resin was washed with DMF-IPA-DMF-IPA-DMF-DCM (x5) sequence with 10 mL of solvent each time. The resin was then dried under vacuum overnight. Loading calibration: To a 4 mL vial was added dried resin (1 mg) and 4-Me-piperidine (20% in DMF, 3 mL). The vial was rocked for 30 min before 100 µL of the liquid was taken for UV measurement. After fitting the UV value to a standard curve, the final loading of resin **8**: L = 1.33 mmol/g, 158 mg.

To a 12 mL syringe with a valve tip was added resin **8** (Loading = 1.33 mmol/g, 0.05 mmol) and 4-Me-piperidine (20% in DMF, 2.5 mL). The syringe was rocked for 5 min at RT twice before filtering out the liquid through the valve. After washing the resin with 5 mL DMF for three time, Fmoc-protected amino acid (0.1 mmol), HATU (0.1 mmol, 38 mg), DMF (2.5 mL), and DIPEA (0.2 mmol, 35 µL) were added to the syringe sequentially. The syringe was rocked for 2 hours at RT before filtering the liquid out through the valve. After acetaldehyde/chloranil test showed the reaction was finished, the resin was washed with DMF-IPA-DMF-IPA-DMF-DCM(x5) sequence with 5 mL of solvent each time. The above procedure was repeated to install each amino acid building blocks. After the installation of the last Fmoc-protected amino acid, the resin was mixed with 4-Me-piperidine (20% in DMF, 2.5 mL). The syringe was rocked for 5 min twice before filtering the liquid out through the valve. 5 mL DMF was used to wash the resin for three time followed by 5 mL DCM wash for three times. After washing, 10 mL 20% HFIP in DCM was added into the syringe, and the syringe was rocked at RT for 15 min. The liquid that contained heptapeptide **SI-4c** was collected and concentrated. The crude product was used in the next macrocyclization step without further purification.

Macrocyclization reaction was carried out under a pseudo-high dilution protocol using two syringe pumps carrying (1) the crude heptapeptide **SI-4c** and DIPEA (0.15 mmol, 26.5 μL) in DMF (2.5 mL), and (2) HATU (0.15 mmol, 57 mg) in DMF (1.5 mL). These two reagents were added at 1.8 mL/h to a flask containing DIPEA (0.15 mmol, 26.5 μL) in DMF (3 mL). After addition, the reaction was allowed to stir at RT and monitored by LC-MS. After full conversion of **SI-4c**, the solvent was removed under reduced pressure. The resulting mixture was dissolved in EtOAc, washed with 1 M HCl, saturated aqueous $NaHCO_3$, and brine. The organic phase was collected and dried over $Na_2SO_4$, dried over $Na_2SO_4$, and concentrated *in vacuo*. CombiFlash chromatography (0-30% Acetone in Hexane) was used to afford **SI-4d** as the product.

**Compound characterization**

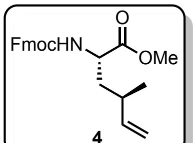

**4**

**4**: (2*S*,4*R*)-2-({[(9H-fluoren-9-yl)methoxy]carbonyl}amino)-4-methylhex-5-enoate.

HRMS (ESI) for $C_{23}H_{25}NNaO_4$ $[M+Na]^+$, 402.1676 (Calc.), found 402.1716.

$^1$H NMR (400 MHz, $CDCl_3$) δ 7.74 (d, *J* = 8 Hz, 2H), 7.58 (t, *J* = 8.0 Hz, 2H), 7.38 (t, *J* = 8Hz, 2H), 7.29 (t, *J* = 8 Hz, 2H), 5.70 – 5.61 (m, 1H), 5.32 (d, *J* = 8 Hz, 1H), 5.05 – 5.00 (m, 2H), 4.41 – 4.35 (m, 3H), 4.21 (t, *J* = 8 Hz, 1H), 3.71 (s, 3H), 2.31 – 2.20 (m, 1H), 1.82 – 1.75 (m, 1H), 1.65 – 1.55 (m, 1H), 1.02 (d, *J* = 4 Hz, 3H).

$^{13}$C NMR (100 MHz, $CDCl_3$) δ 173.5, 156.0, 144.1, 143.9, 142.6, 141.4, 141.4, 127.8, 127.1, 125.2, 125.2, 120.17, 114.9, 67.0, 52.6, 52.4, 47.3, 39.4, 34.9, 20.9.

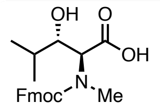

**SI-2g**

**SI-2g**: (2*S*,3*S*)-2-{[(9H-fluoren-9-ylmethoxy)carbonyl](methyl)amino}-3-hydroxy-4-methylpentanoic acid

HRMS (ESI) for $C_{22}H_{24}NO_5$ $[M-H]^-$, 382.1660 (Calc.), found 382.1658.

$^1$H NMR (400 MHz, MeOD) δ 7.78 – 7.75 (m, 2H), 7.64 – 7.57 (m, 2H), 7.39 – 7.35 (m, 2H), 7.33 – 7.26 (m, 2H), 4.57 – 4.29 (m, 3H), 4.23 – 4.17 (m, 1H), 3.81 – 3.78 (m, 1H), 2.89 (s, 1H), 2.78 (s, 2H), 1.71 - 1.60 (m, 1H), 0.97 (d, *J* = 4 Hz, 2H), 0.88 – 0.84 (m, 3H), 0.77 (d, *J* = 4 Hz, 1H). This compound has multiple rotamers; only major peaks are listed and integrated.

$^{13}$C NMR (100 MHz, MeOD) δ 174.1, 173.8, 158.3, 158.0, 145.4, 145.4, 145.2, 145.2, 142.8, 142.8, 142.7, 142.7, 128.9, 128.9, 128.3, 128.3, 126.3, 126.1, 126.1, 126.1, 121.1, 121.1, 121.1, 75.3, 75.1, 69.1, 68.8, 62.2, 48.7, 37.9, 35.3, 32.5, 32.4, 31.0, 30.8, 20.5, 20.4, 16.2, 15.7. This compound has multiple rotamers; only major peaks are listed.

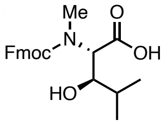

**SI-3h**

**SI-3h**: (2*S*,3*R*)-2-{[(9H-fluoren-9-ylmethoxy)carbonyl](methyl)amino}-3-hydroxy-4-methylpentanoic acid

HRMS (ESI) for $C_{22}H_{24}NO_5$ $[M-H]^-$, 382.1660 (Calc.), found 382.1658.

$^1$H NMR (400 MHz, MeOD) δ 7.80 – 7.76 (m, 2H), 7.64 – 7.58 (m, 2H), 7.40 – 7.35 (m, 2H), 7.33 – 7.26 (m, 2H), 4.84 (d, *J* = 4 Hz, 1H), 4.56 – 4.37 (m, 2H), 4.28 – 4.19 (m, 2H), 3.86 – 3.82 (m, 1H), 3.50 – 3.45 (m, 1H), 3.00 (s, 3H), 1.66 – 1.55 (m, 1H), 1.17 (t, *J* = 8 Hz, 1H), 1.01 (d, *J* = 4 Hz, 2H), 0.93 (d, *J* = 4 Hz, 1H), 0.87 (d, *J* = 4 Hz, 2H), 0.70 (d, *J* = 8 Hz, 1H). This compound has multiple rotamers; only major peaks are listed and integrated.

$^{13}$C NMR (100 MHz, MeOD) δ 173.9, 173.6, 159.3, 158.6, 145.4, 145.3, 145.3, 142.8, 142.8, 142.7, 142.7, 128.9, 128.9, 128.3, 128.3, 128.2, 126.1, 126.0, 126.0, 121.0, 77.5, 76.3, 69.0, 69.0, 67.0, 62.6, 62.1, 48.5, 33.8, 33.0, 32.6, 32.3, 26.4, 20.0, 19.9, 19.2, 18.4, 15.5. This compound has multiple rotamers; only major peaks are listed.

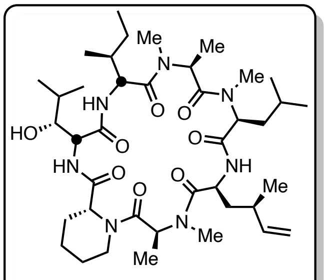

**Ternatin-4**

**Ternatin-4**, 0.05 mmol scale, 70% overall yield, 27 mg, white solid.

Spectral data of **ternatin-4** are in accordance with our previous report.[8]

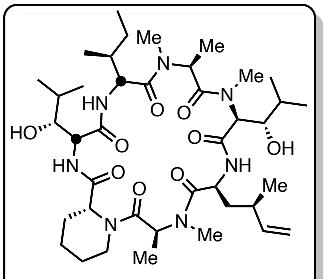

**(*S,S*)-A3**

**(*S,S*)-A3**, 0.03 mmol scale, 21% overall yield, 5 mg, white solid.

HRMS (ESI) for $C_{40}H_{69}N_7NaO_9$ $[M+Na]^+$, 814.5049 (Calc.), found 814.5059.

$^1$H NMR (400 MHz, Acetone-d$^6$) δ 7.83 – 7.67 (m, 2H), 7.45 (m, 1H), 5.75 – 5.52 (m, 1H), 5.39 (m, 1H), 5.18 – 4.77 (m, 3H), 4.74 – 4.51 (m, 1H), 4.44 – 4.24 (m, 1H), 4.19 – 3.85 (m, 1H), 3.60 – 3.35 (m, 2H), 3.07 (d, *J* = 5.6 Hz, 1H), 3.02 – 2.88 (m, 9H), 2.43 – 2.30 (m, 1H), 2.17 (m, 2H), 1.84 – 1.69 (m, 6H), 1.53 – 1.39 (m, 7H), 1.32 – 1.22 (m, 5H), 0.99 – 0.91 (m, 10H), 0.88 – 0.72 (m, 16H). This compound has multiple rotamers; only major peaks are listed and integrated.

$^{13}$C NMR (100 MHz, Acetone-d$^6$): δ 174.5, 174.4, 174.3, 174.0, 172.9, 170.2, 169.2, 143.5, 115.6, 80.8, 76.2, 72.9, 64.6, 56.3, 56.1, 53.0, 52.1, 50.4, 50.2, 44.2, 40.71, 36.7, 34.8, 34.18, 31.8, 27.2, 26.5, 26.3, 26.10, 25.96, 25.67, 23.33, 21.81, 21.48, 21.25, 21.06, 21.03, 20.73, 18.28, 15.70, 15.66, 15.60, 15.52, 21.9, 21.5, 15.5, 14.8, 14.4, 14.1, 12.2, 12.1. This compound has multiple rotamers; only major peaks are listed and integrated.

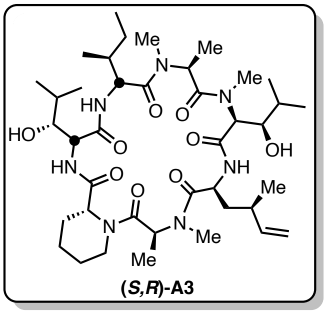

**(*S,R*)-A3**, 0.1 mmol scale, 35% overall yield, 27.2 mg, white solid.

HRMS (ESI) for $C_{40}H_{69}N_7NaO_9$ $[M+Na]^+$, 814.5049 (Calc.), found 814.5059.

$^1$H NMR (400 MHz, Acetone-d$^6$) δ 7.78 – 7.75 (m, 2H), 7.52 (d, J = 8 Hz, 1H), 5.85 (q, *J* = 8 Hz, 1H), 5.76 – 5.67 (m, 1H), 5.18 (br, 1H), 5.08 – 4.95 (m, 3H), 4.90 – 4.85 (m, 1H), 4.80 – 4.72 (m, 2H), 4.09 – 4.02 (m, 2H), 3.92 – 3.89 (m, 1H), 3.63 – 3.59 (m, 1H), 3.10 (s, 3H), 3.05 (s, 3H), 2.98 (s, 3H), 2.52 – 2.49 (m, 1H), 2.22 – 2.14 (m, 1H), 2.00 – 1.72 (m, 6H), 1.65 – 1.56 (m, 4H), 1.53 – 1.44 (m, 2H), 1.40 – 1.35 (m, 3H), 1.33 – 1.26 (m, 3H), 1.13 (d, *J* = 8 Hz, 3H), 1.05 – 0.94 (m, 18H), 0.90 (d, *J* = 8 Hz, 3H). This compound has multiple rotamers; only major peaks are listed and integrated.

$^{13}$C NMR (100 MHz, Acetone-d$^6$) δ 174.6, 174.5, 174.4, 173.3, 171.7, 169.2, 168.7, 143.6, 115.5, 76.1, 71.8, 63.8, 56.3, 56.1, 53.1, 51.8, 50.9, 49.7, 44.3, 36.7, 34.6, 31.8, 27.4, 26.6, 26.2, 21.9, 21.5, 21.5, 21.2, 15.9, 15.5, 14.7, 14.2, 14.1, 12.1. This compound has multiple rotamers; only major peaks are listed.

**NMR Spectra:**

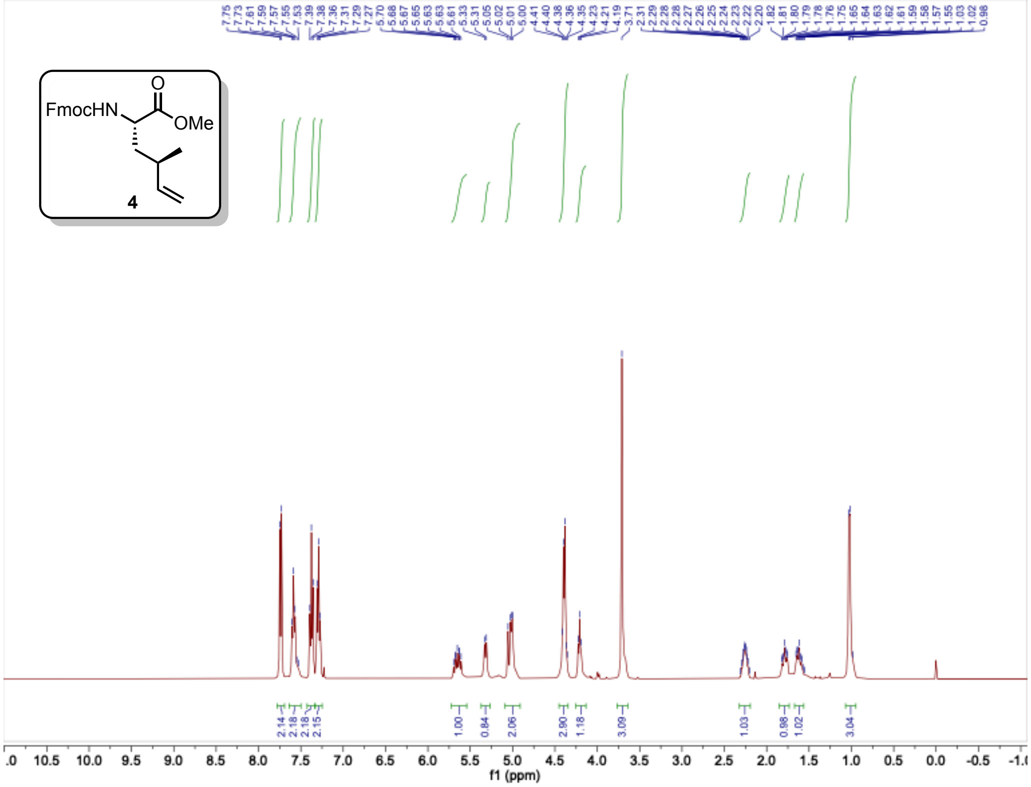
FmocHN
O
OMe
4
f1 (ppm)

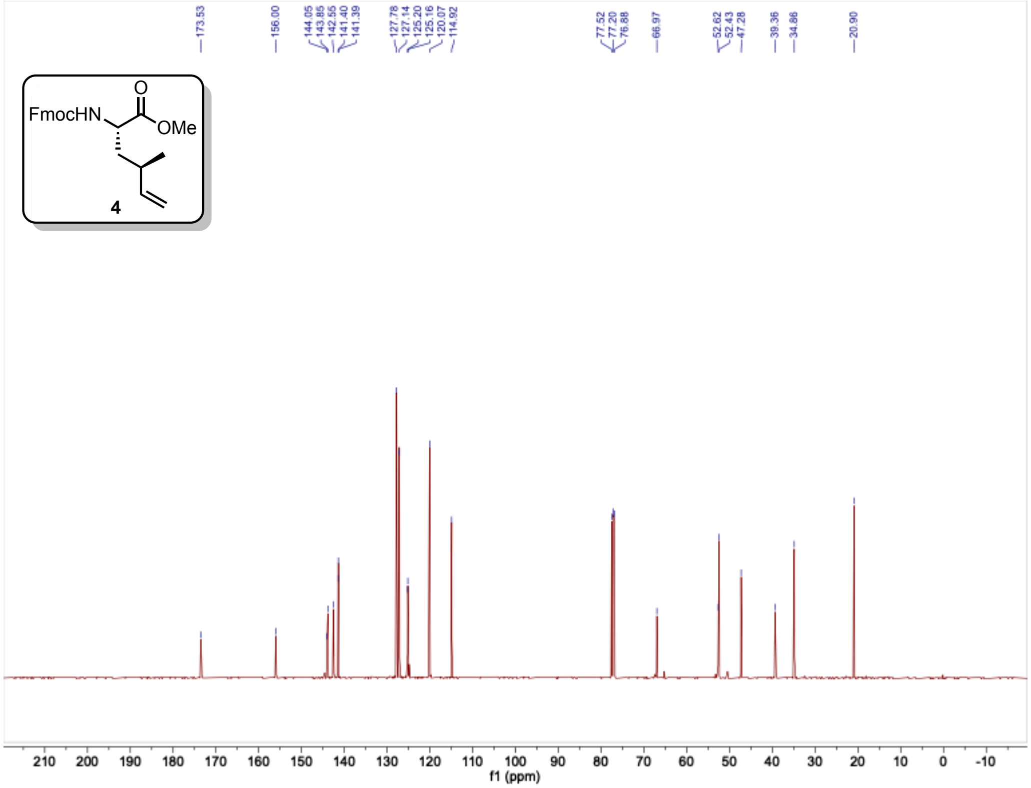
FmocHN
O
OMe
4
173.53
156.00
144.05
143.85
142.55
141.40
141.39
127.78
127.14
125.20
125.16
120.07
114.92
77.52
77.20
76.88
66.97
52.62
52.43
47.28
39.36
34.86
20.90
f1 (ppm)

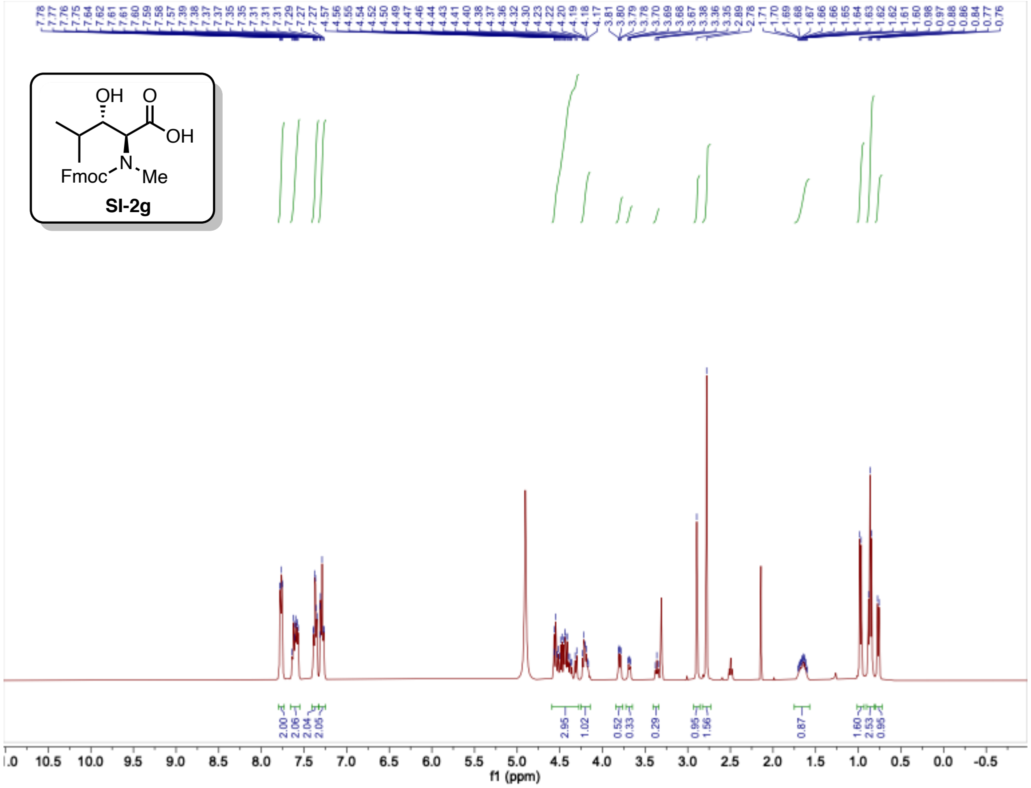
SI-2g
f1 (ppm)

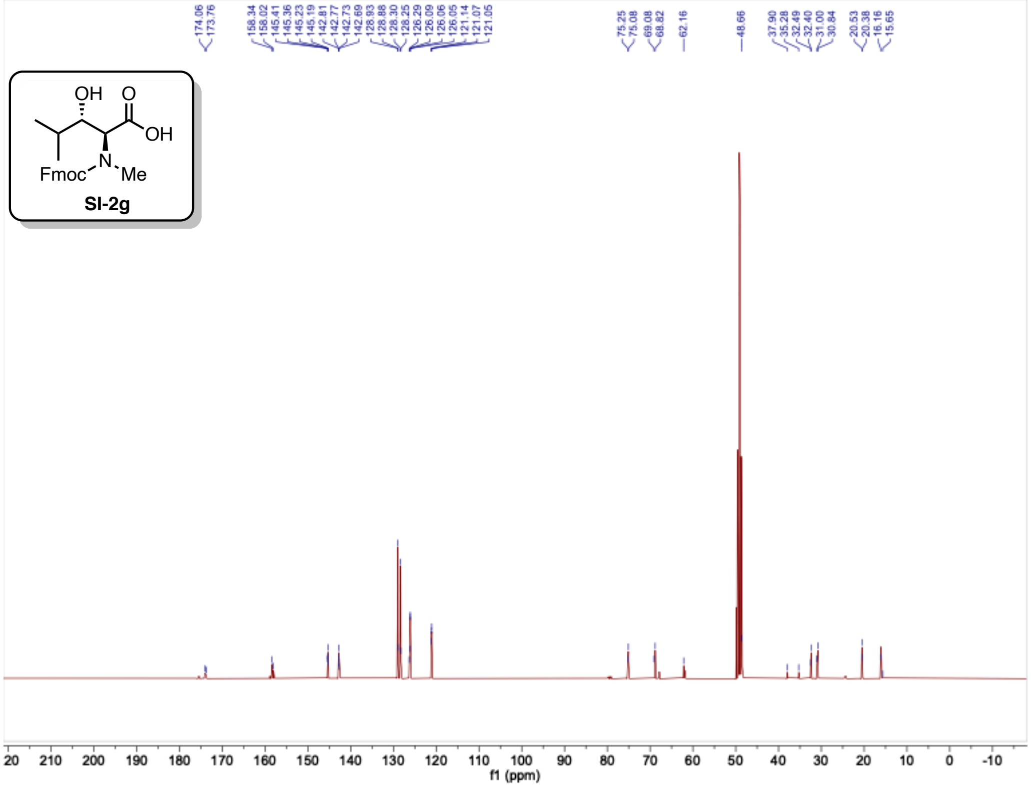
SI-2g
f1 (ppm)

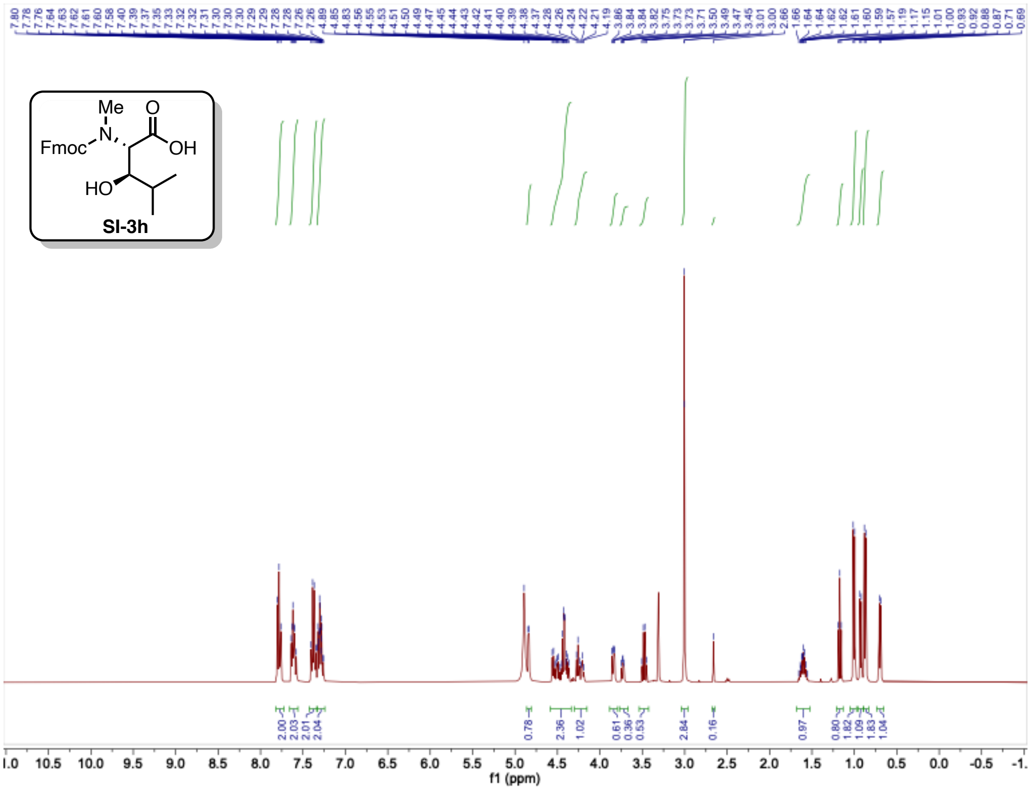
Me
O
Fmoc
N
OH
HO
SI-3h
f1 (ppm)

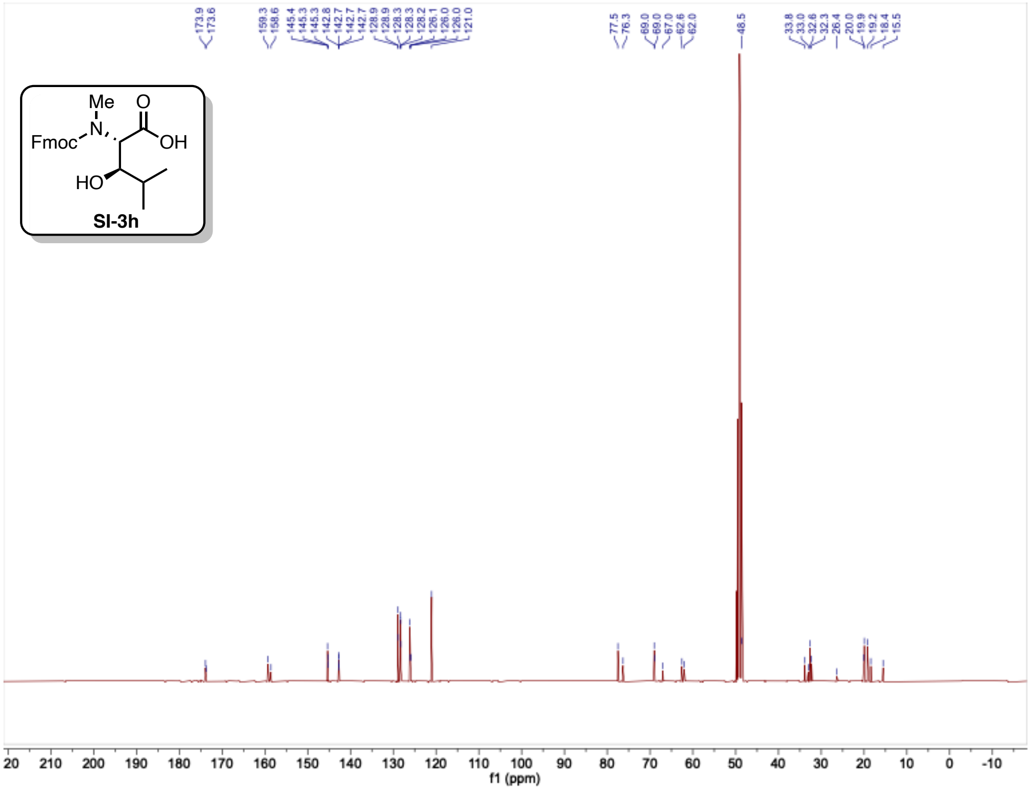
Me
O
Fmoc
N
OH
HO
SI-3h
f1 (ppm)

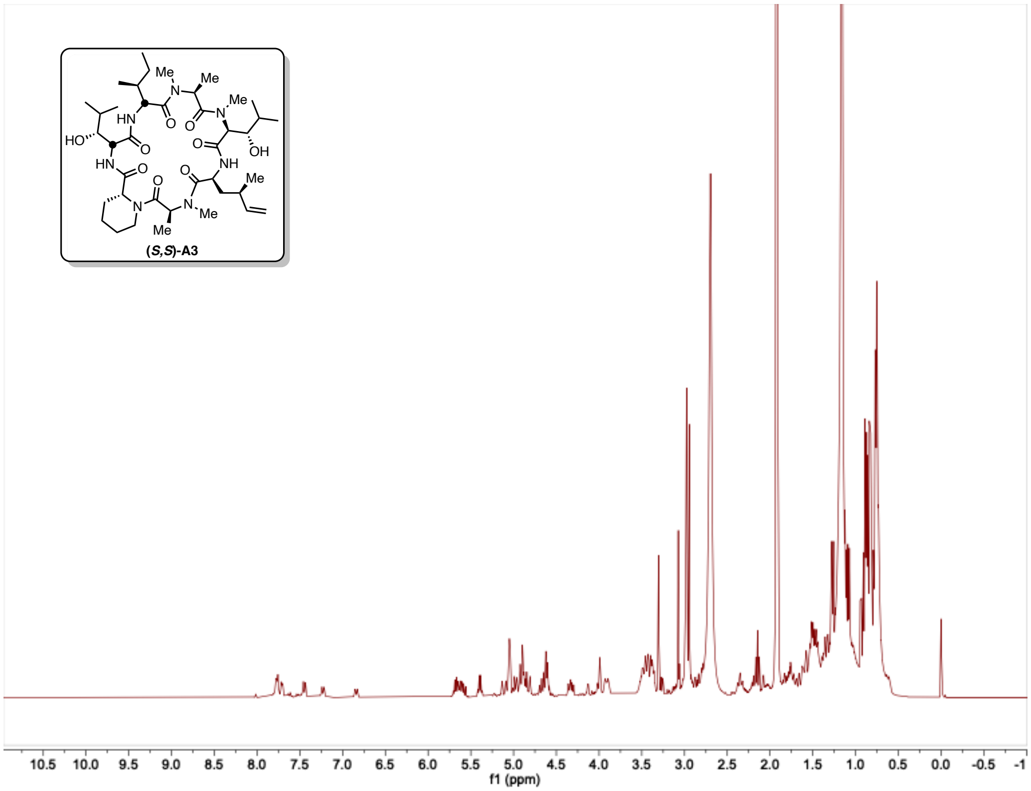
(S,S)-A3
f1 (ppm)

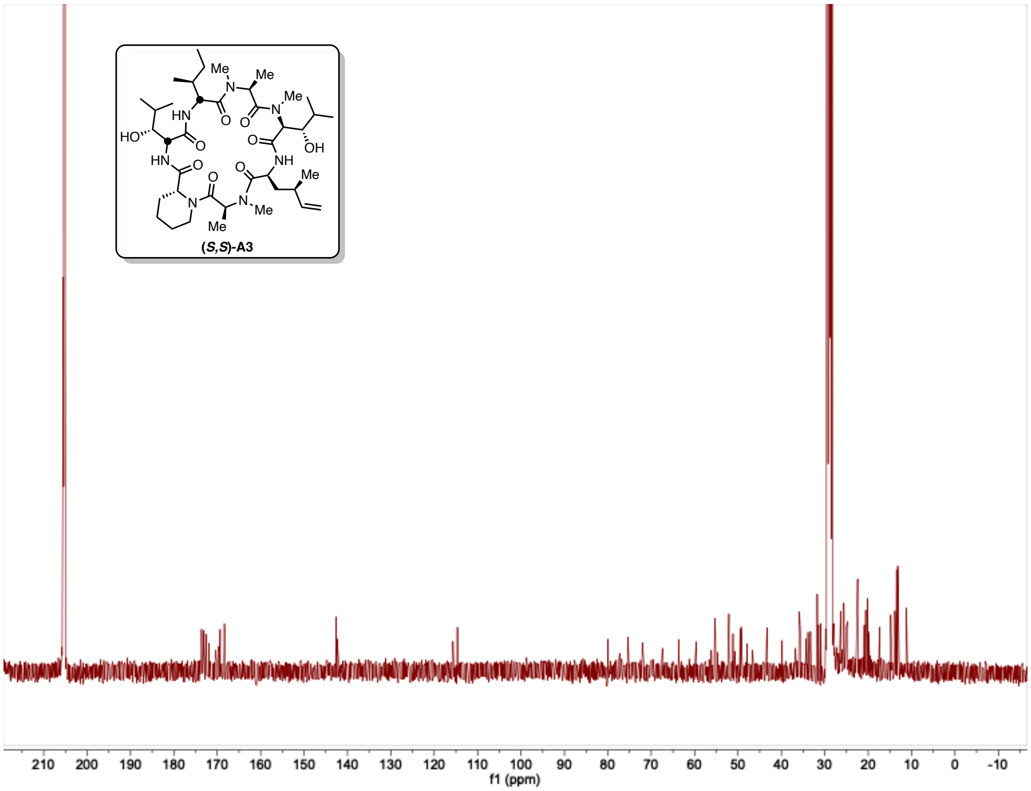
(S,S)-A3
f1 (ppm)

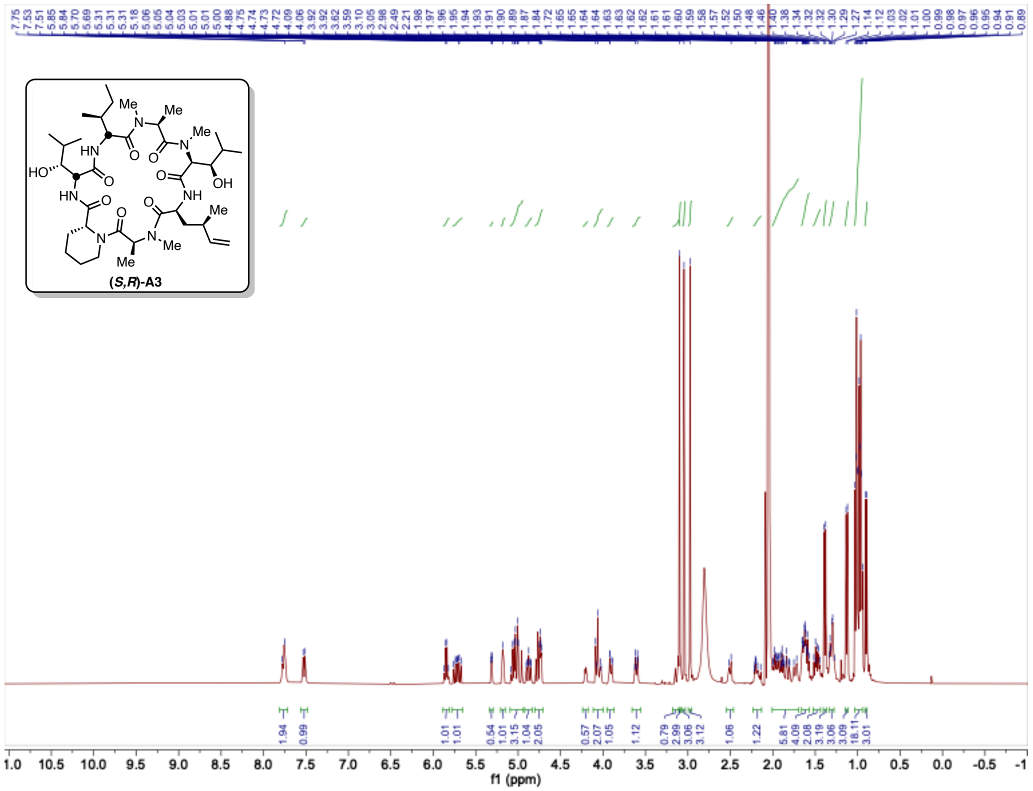

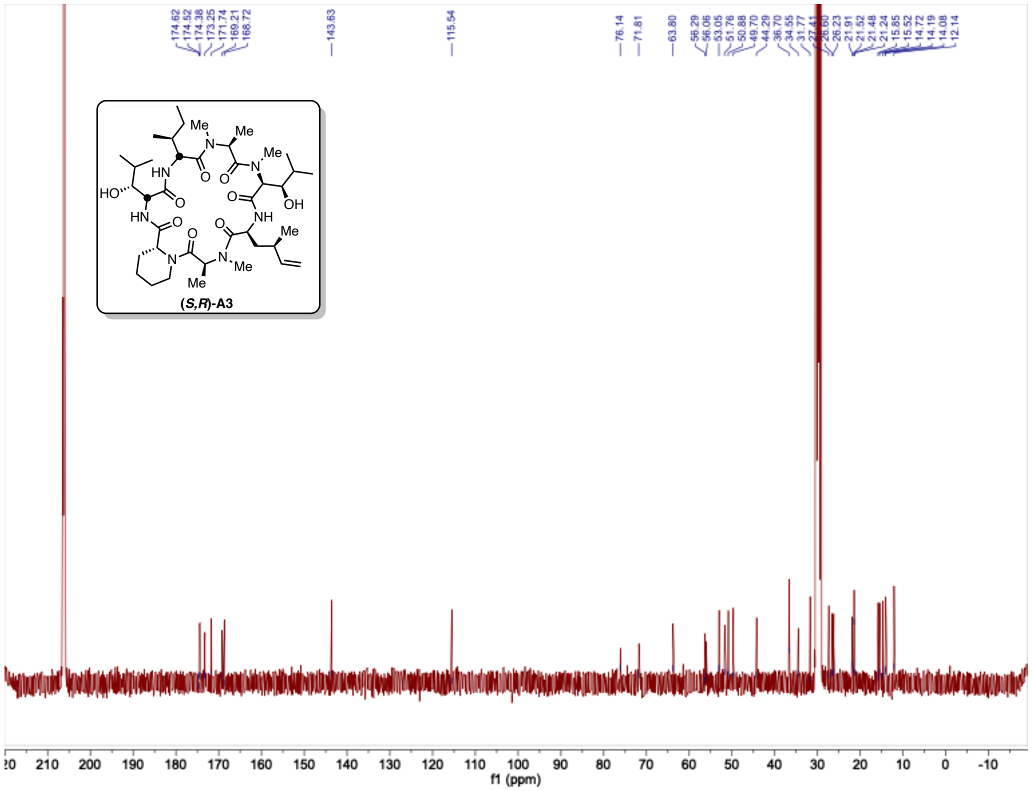

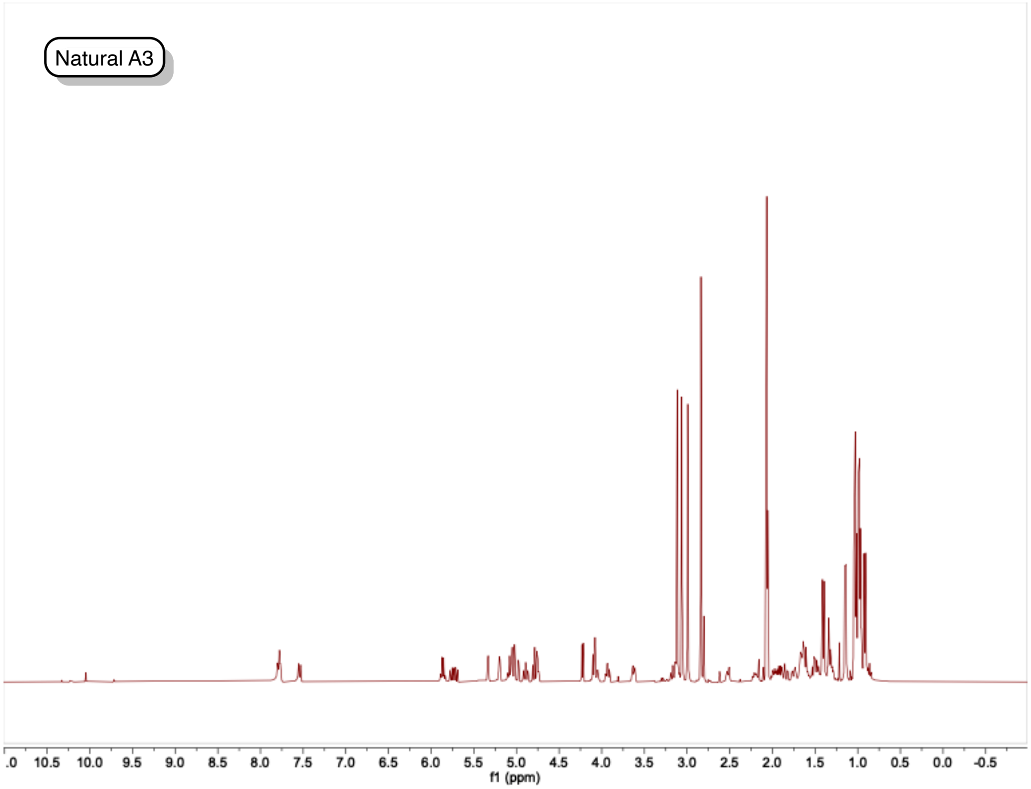
Natural A3
.0
10.5
10.0
9.5
9.0
8.5
8.0
7.5
7.0
6.5
6.0
5.5
5.0
4.5
4.0
3.5
3.0
2.5
2.0
1.5
1.0
0.5
0.0
-0.5
f1 (ppm)

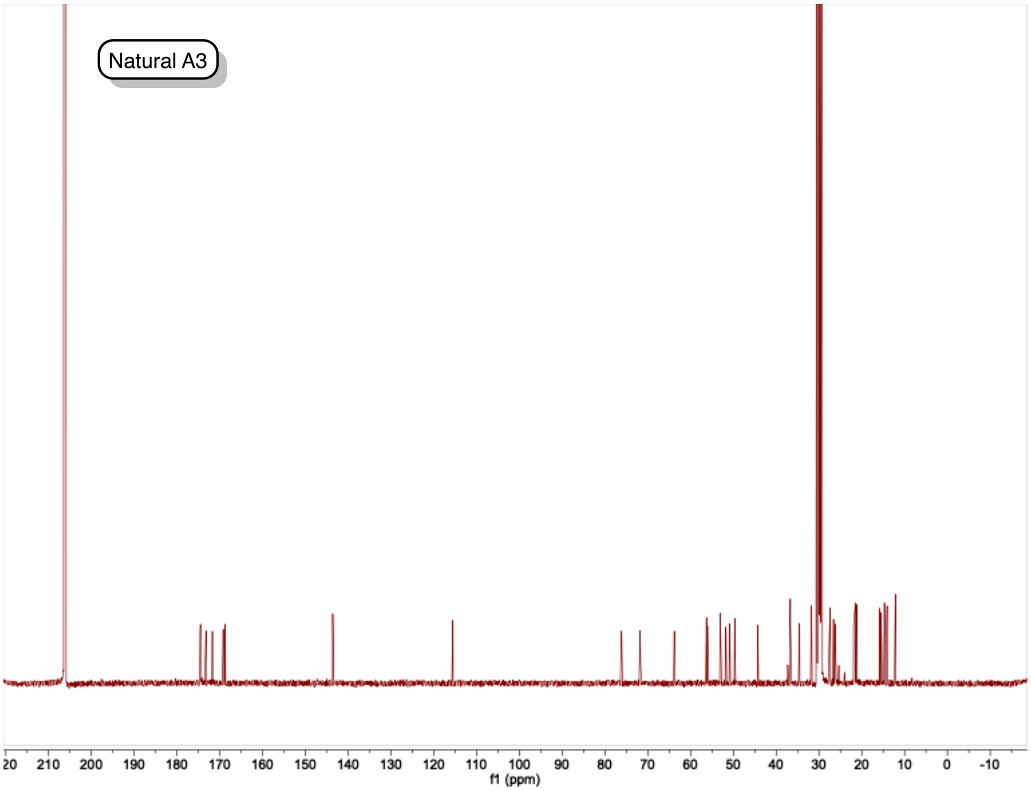
Natural A3
20
210
200
190
180
170
160
150
140
130
120
110
100
90
80
70
60
50
40
30
20
10
0
-10
f1 (ppm)
